# STRAIGHT-IN: A platform for high-throughput targeting of large DNA payloads into human pluripotent stem cells

**DOI:** 10.1101/2021.12.08.471715

**Authors:** Catarina Grandela, Albert Blanch-Asensio, Karina O. Brandão, Tessa de Korte, Loukia Yiangou, Mervyn P.H. Mol, Berend J. van Meer, Christine L. Mummery, Richard P. Davis

**Author notes:** These authors contributed equally (Catarina Grandela, Albert Blanch-Asensio, Karina O. Brandão, Tessa de Korte). Correspondence (R.P.D).

## Abstract

Inserting large DNA payloads (>10 kb) into specific genomic sites of mammalian cells remains challenging. Applications ranging from synthetic biology to evaluating the pathogenicity of disease-associated variants for precision medicine initiatives would greatly benefit from tools that facilitate this process. Here, we merge the strengths of different classes of site-specific recombinases and combine these with CRISPR/Cas9-mediated homologous recombination to develop a strategy for stringent site-specific replacement of genomic fragments at least 50 kb in size in human induced pluripotent stem cells (hiPSCs). We demonstrate the versatility of STRAIGHT-IN (<u>S</u>erine and <u>T</u>yrosine <u>R</u>ecombinase <u>A</u>ssisted Integration of <u>G</u>enes for <u>H</u>igh-<u>T</u>hroughput <u>IN</u>vestigation) by: (i) inserting various combinations of fluorescent reporters into hiPSCs to assess excitation-contraction coupling cascade in derivative cardiomyocytes, and; (ii) simultaneously targeting multiple variants associated with inherited cardiac arrhythmic disorder into a pool of hiPSCs. STRAIGHT-IN offers a precise approach to generate genetically-matched panels of hiPSC lines efficiently and cost-effectively.

## INTRODUCTION

Human pluripotent stem cells (hPSCs) have tremendous potential for advancing the understanding of human development and disease. To realise this, methods for efficient targeted genetic modification are crucial. Endonuclease-based gene editing systems (i.e. CRISPR-Cas9) have made it significantly easier to perform small-scale genomic modifications in hPSCs. However, strategies to insert multi-kilobase (kb) payloads that consist of several transgenes or a genomic fragment, for example, are limited since the efficiency of homology directed repair-mediated targeting decreases significantly with increasing insert size (Byrne et al., 2015; Zhang et al., 2021).

Site-specific recombinases (SSRs) are useful tools for performing difficult genome engineering tasks, (e.g. insertion, deletion or inversion of large DNA segments), in cultured human cells (Turan et al., 2013). SSRs are divided into two families based on the identity of the nucleophilic active site amino acid residue. Phage-derived serine recombinases, which include Bxb1 and φC31 integrases, mediate unidirectional recombination between the enzyme’s unique phage and bacterial attachment sites (*attP* and *attB* respectively), thereby enabling irreversible integration of transgenes into the genome (Brown et al., 2011). For instance, it has been demonstrated that payloads of up to 33 kb can be efficiently inserted using Bxb1 in Chinese hamster ovary (CHO) cells (Gaidukov et al., 2018). Tyrosine recombinases (e.g. Cre or FLP recombinases) can also integrate DNA fragments, although their integration efficiencies are relatively lower and the event reversible (Turan et al., 2013). However, they are effective at excising cassettes flanked by directly repeated recombination sites, leading to their widespread use in conditionally knocking out genes, removing selection cassettes following targeting and in lineage tracing studies (Davis et al., 2008; Skarnes et al., 2011).

Despite both classes of SSRs being used to genetically modify hPSCs (Chaudhari et al., 2020; Du et al., 2009; Ordovás et al., 2015; Zhu et al., 2014), efficient methods to perform targeted integrations of large DNA payloads (>10 kb) that are also suitable for multiplex assays are still lacking. Cassette exchange strategies, whereby a previously targeted ‘landing pad’ (LP) cassette containing SSR recognition/attachment sites is used to insert transgenes flanked by corresponding sequences, have been developed for repeated modifications of hPSCs or differentiated progenitor cells (Lv et al., 2018; Pei et al., 2015; Zhu et al., 2014). However, the largest payload reported to be inserted was ∼7 kb (Zhu et al., 2014). Alternatively, φC31 and λ integrases can introduce larger inserts (up to ∼20 kb) but only into pseudo-*attP* sites already present in the human genome, thus preventing selection of the target site (Chaudhari et al., 2020; Farruggio et al., 2017; Liu et al., 2009). Furthermore, locus-specific silencing of the transgenes occurred in some instances (Farruggio et al., 2017).

Here we have coalesced the advantages of both classes of SSRs to develop a platform, which we term STRAIGHT-IN (for <u>S</u>erine and <u>T</u>yrosine <u>R</u>ecombinase <u>A</u>ssisted Integration of <u>G</u>enes for <u>H</u>igh-<u>T</u>hroughput <u>IN</u>vestigation). STRAIGHT-IN enables the targeted integration or substitution of large genomic fragments into hiPSCs while leaving minimal traces (<300 bp) of plasmid backbone DNA sequences in the locus. We demonstrate how the platform can be applied to construct synthetic genetic circuits, by generating a cell line containing multiple genetic reporters (∼14 kb payload) to assess excitation-contraction coupling in hiPSC-derived cardiomyocytes (hiPSC-CMs). Furthermore, we showcase the ability of STRAIGHT-IN to support multiplex genetic assays by simultaneously generating a library of genetically-matched hiPSC lines carrying heterozygous mutations in the gene *KCNH2*, which can result in various cardiac arrhythmia syndromes in patients (Chen et al., 2016). We confirm that the hiPSC-CMs for the *KCNH2* variant A561T, reflected the expected electrophysiological disease phenotype. Overall, these results highlight the possibilities offered by STRAIGHT-IN for expanding the range of biological questions that can be investigated using hiPSCs in a high-throughput manner.

## RESULTS

### hiPSC acceptor lines for Bxb1- and φC31-mediated integration

For targeted integration of large genomic fragments into hiPSCs, we first generated acceptor lines that contained a LP cassette. The LP cassettes for both Bxb1 and φC31 were similarly designed and included a constitutive promoter (phosphoglycerate kinase promoter; pGK) driving expression of a fluorescent protein, an *attP* site recognised by the corresponding serine recombinase, and an antibiotic positive selection marker without an ATG initiation codon. These cassettes were also flanked by heterotypic recognition sites for either Cre or Flp recombinase to enable their excision downstream.

The *AAVS1* locus can support stable, long-term expression of introduced transgenes, including in differentiated hiPSC derivatives such as hiPSC-CMs (Sun et al., 2020). We initially targeted a single copy of each LP cassette to *AAVS1*, generating the hiPSC acceptor lines *AAVS*1-Bxb1 and *AAVS1*-φC31 (**Figure 1A**). Since each LP construct was comprised of unique components, we also generated an hiPSC line that was biallelically targeted with both LPs (*AAVS1*-Dual), thereby providing a cell line containing an orthogonal pair of target sites that do not cross-react. Fluorescent reporters facilitated the isolation of clonal hiPSCs expressing either GFP (*AAVS1*-Bxb1), mCherry (*AAVS1*-φC31), or both in the case of the *AAVS1*-Dual acceptor line (**Figure 1B**). Screening and genotyping PCR confirmed the LP cassettes were correctly targeted to the *AAVS1* locus for each of the hiPSC acceptor lines (**Figures 1C, S1A and S1B**). In addition, ddPCR for the genes *eGFP, mCherry, BsdR* and *BleoR* verified only a single integration of each LP in the respective acceptor lines (**Figure 1D**).

**Figure 1.**
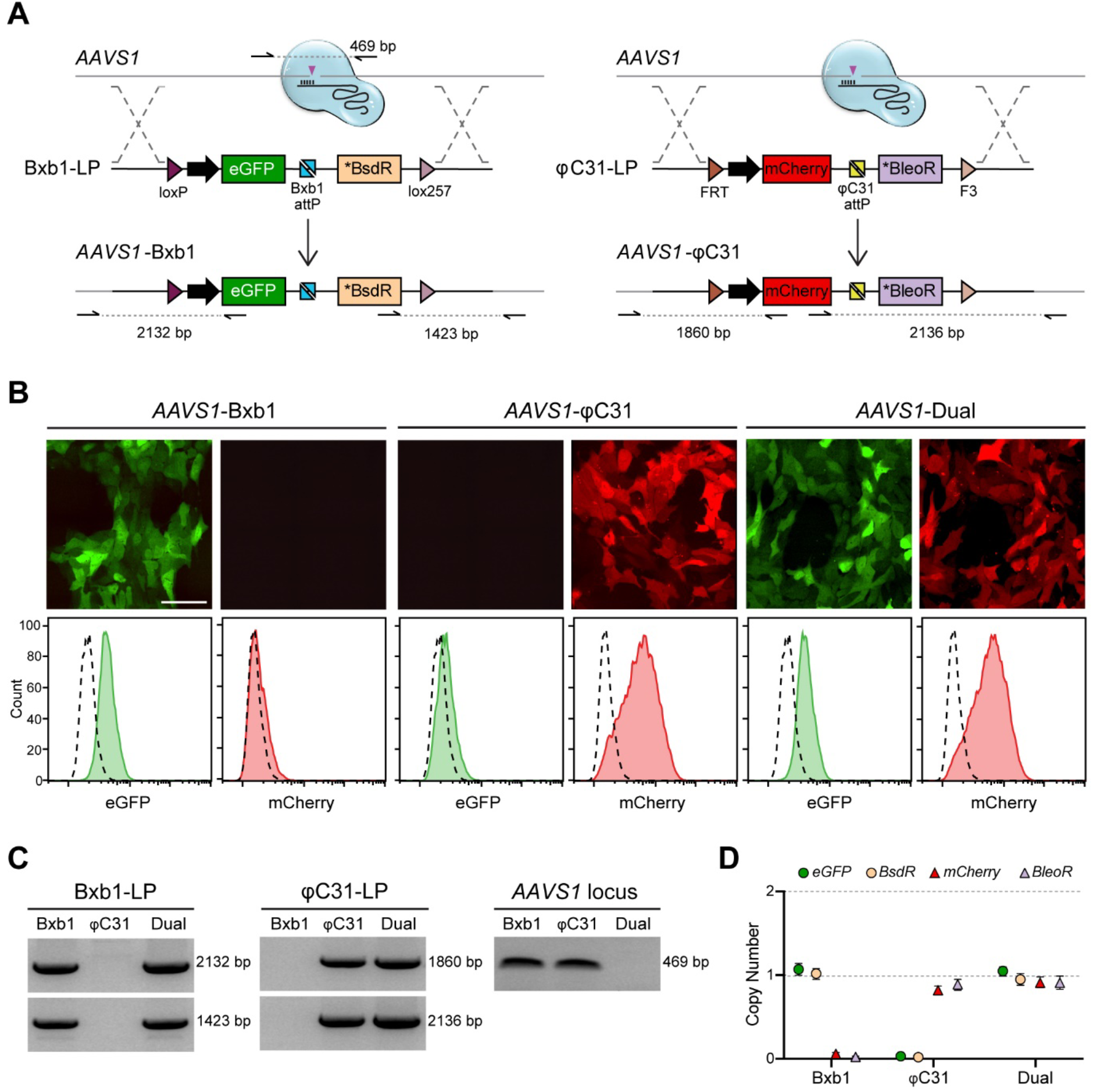
hiPSC acceptor lines for Bxb1- and φC31-mediated integration. (A) Schematic for the targeting of the Bxb1- and φC3-landing pad (LP) cassettes to the *PPP1R12C* locus (*AAVS1*). The large black arrows represent the pGK promoter driving constitutive expression of the fluorescent reporter genes, eGFP or mCherry. The * indicates that the antibiotic positive selection marker (blasticidin S deaminase (BsdR) or bleomycin resistance (BleoR)) lacks an ATG initiation codon and so is not expressed in the resulting targeted hiPSCs. Purple triangles indicate the sites of CRISPR/Cas9-induced double-strand DNA breaks. Half arrows indicate primer binding sites with dotted lines representing the resulting PCR amplicons. (B) Fluorescence images (*top*) and flow cytometry plots (*below*) indicating the expression of the eGFP and mCherry reporters in the resulting targeted *AAVS1*-acceptor hiPSC lines. Scale bar, 50 µm. (C) PCR amplification, using primer pairs indicated in panel (A), of genomic DNA confirming targeting of the LP cassettes in the resulting *AAVS1*-acceptor hiPSC lines. The LP cassette was targeted to only one allele in the *AAVS1*-Bxb1 and *AAVS1*-φC31 hiPSCs, while both alleles were targeted in the *AAVS1*- Dual hiPSCs (*right panel*). (D) ddPCR confirming the *AAVS1*-acceptor hiPSC lines had either 0 or 1 copy of each LP cassette inserted into the genomic DNA. Error bars represent Poisson 95% CI.

### Bxb1 mediates effective integration of DNA into hiPSCs without size restrictions

To evaluate the efficiency and specificity of the system for exogenous DNA integration, donor vectors specific for both Bxb1- and φC31-LPs were developed (**Figures S1C-E**) and co-transfected with a plasmid expressing either Bxb1 or φC31 into *AAVS1*-Dual. These donor vectors consisted of a constitutive promoter (elongation factor-1 alpha; EF1a) preceding an ATG initiation codon and the *attB* sequence recognised by either Bxb1 or φC31. Correct integration of the construct into the respective LP would form two new recombination sites (*attR* and *attL*), as well as place the promoter and start codon in the donor vector upstream and in frame with the antibiotic positive selection marker, thereby enabling enrichment of correctly integrated clones by either blasticidin (Bxb1-LP) or zeocin (φC31-LP) selection (**Figure 2A**).

**Figure 2.**
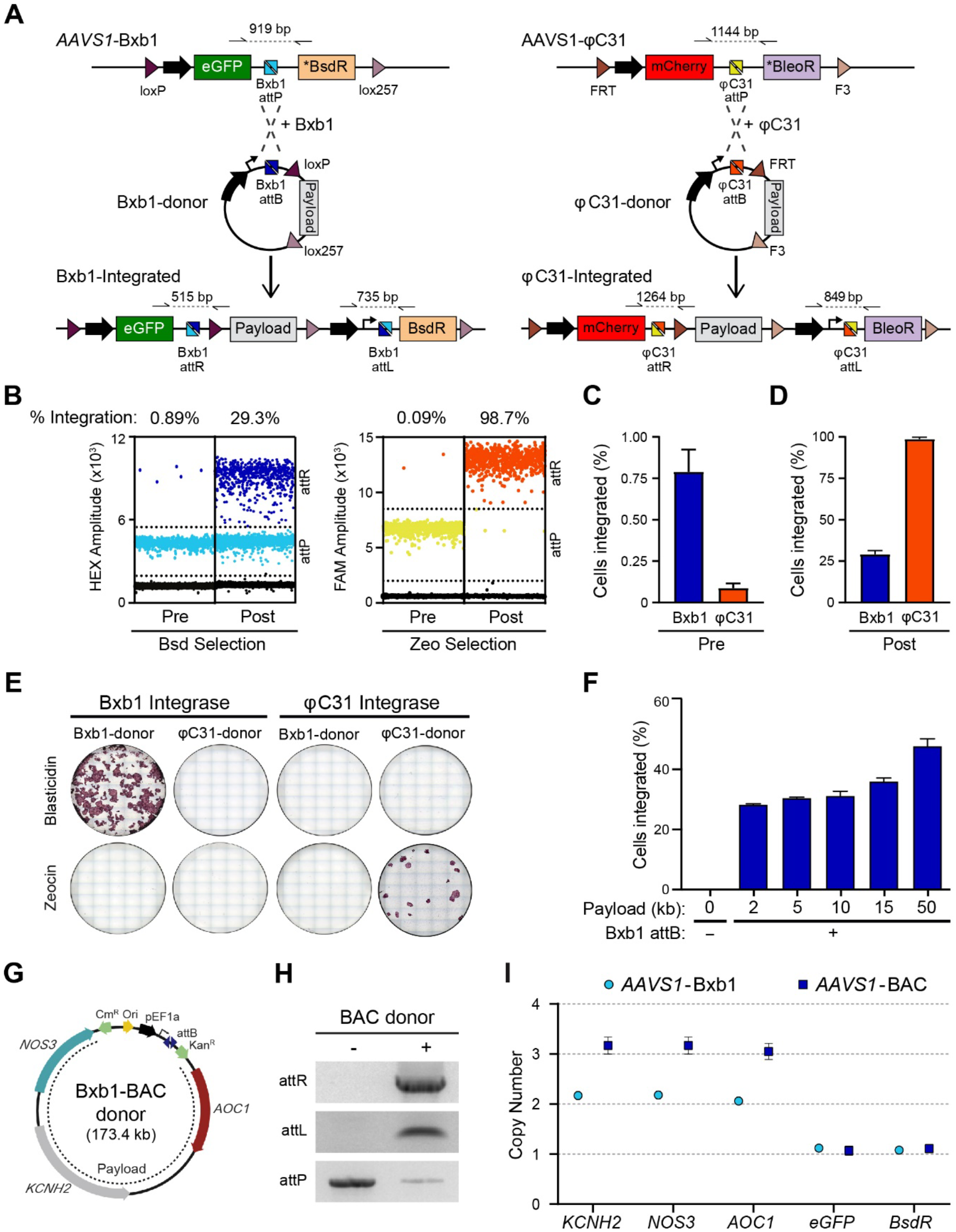
Targeted integration of DNA payloads into hiPSC acceptor lines. (A) Schematic for integrating donor constructs by either Bxb1 (*left*) or φC31 (*right*) integrases into the corresponding LP cassette targeted to *AAVS1*. Correct integration of the donor construct results in expression of the antibiotic positive selection marker present in the LP cassette. The large black arrows represent constitutive promoters, while “Payload” indicates the region in the donor construct where DNA sequences to integrate are inserted. Half arrows indicate primer binding sites with dotted lines representing the resulting PCR amplicons. (B) Representative ddPCR dot plots indicating the integration of either the empty Bxb1 (dark blue) or φC3- (orange) donor vector into *AAVS1-*Dual hiPSCs both before (Pre) and after (Post) antibiotic enrichment. Dots represent droplets containing the indicated sequence (attR or attP), while the percentages denote the calculated integration efficiency. (C, D) Calculated mean integration efficiency of the Bxb1- and φC3-donor vectors both before (C) and after (D) antibiotic enrichment from 3 transfections. Error bars represent ± SEM. (E) Alkaline phosphatase staining of hiPSCs following enrichment of cells transfected with plasmids expressing either Bxb1 or φC31 integrase as well as the Bxb1- or φC31-donor vectors demonstrating that integration is both integrase- and selection cassette-specific. (F) Average percentage of hiPSCs that integrated donor constructs with or without the Bxb1-*attB* sequence (+ or -) and payloads ranging from 0 – 50 kb into the Bxb1-LP following one round of blasticidin selection. n=4 transfections; error bars represent ± SEM. (G) Schematic of the Bxb1-BAC donor vector indicating the 3 genes (*KCNH2, NOS3, AOC1*) encoded by the BAC, as well as modifications made to enable targeted integration of the vector. CmR, chloramphenicol acetyltransferase. (H) PCR amplification, using primer pairs indicated in (A), of genomic DNA confirming integration of the Bxb1-BAC donor vector into a subset of the *AAVS1-*Bxb1 hiPSCs. The “-” and “+” symbols indicate before and after blasticidin selection respectively. (I) ddPCR confirming that a hiPSC clone (*AAVS1*-BAC) identified as having the Bxb1-BAC donor vector integrated into the Bxb1-LP contained 3 copies of *KCNH2, NOS3* and *AOC1*. The non-integrated *AAVS1*-Bxb1 hiPSC line retained 2 copies of each. Both lines contained a single copy of the LP cassette transgenes, *eGFP* and *BsdR*. Error bars represent Poisson 95% CI.

To detect and quantify the proportion of cells that had integrated the donor vector, ddPCR assays were developed that distinguished both integrated and non-integrated cells (**Figure S2**). This assay indicated that Bxb1 was ∼10-fold more efficient at mediating the integration of the donor vector than φC31 (**Figures 2B and 2C**). Additionally, only the integration event mediated by Bxb1 could be detected by diagnostic PCR prior to antibiotic selection (**Figure S3A**). Although non-integrated cells were more sensitive to zeocin selection than blasticidin (**Figure 2D**), integration is a more critical step when performing multiplex targeting. Furthermore, enrichment with blasticidin selection could be improved by maintaining the hiPSCs with the antibiotic for a longer period (**Figure S3B**). Therefore, we focused on further evaluating the capabilities of Bxb1 to perform targeted integrations of large DNA payloads into hiPSCs. The specificity of the integrases to their target sequences was also evident, with no colonies obtained when we mismatched the integrase with the *attB* donor vector (**Figure 2E**).

To evaluate whether there was a limit on the size of the donor construct that could be integrated, a series of donor vectors with DNA payloads varying in size from ∼2 to ∼50 kb were co-transfected with Bxb1. Between 30-50% of the cells following enrichment had the payload integrated (**Figures 2F and S3C**), suggesting that overall targeting frequency was independent of size. We did not detect an upper limit to the length of the payload that could be targeted, with a modified 173 kb bacterial artificial chromosome (BAC) construct also readily integrated into the *AAVS1* locus and the resulting hiPSCs acquiring an additional copy of the three genes (*KCNH2, NOS3* and *AOC1*) present on the BAC donor (**Figures 2G-I**).

### Cre and FLP efficiently excise auxiliary sequences following donor vector integration

Upon integration, the entire donor vector is integrated into the LP. These auxiliary sequences within the vector backbone can lead to silencing of either the transgenes or neighbouring genetic elements (Chen et al., 2004; Pham et al., 1996) with the effect prevented or even reversed if these sequences are subsequently excised from the targeted loci (Davis et al., 2008; Riu et al., 2005). Including *loxP* and *lox257* sequences in both the LP construct and donor vector resulted in both the vector backbone and the majority of the LP cassette being flanked by these recombination sequences following vector integration (**Figure 3A**). Transiently expressing Cre led to the excision of these auxiliary sequences in >90% of integrated recombinant clones, leaving only the integrated DNA payload plus a single copy of *loxP* and *lox257* at the targeted *AAVS1* locus (**Figures 3B and 3C**). We also confirmed that FLP recombinase could efficiently excise FRT- and F3-flanked sequences following φC31-mediated integration of donor vectors (**Figures S3D-F**).

**Figure 3.**
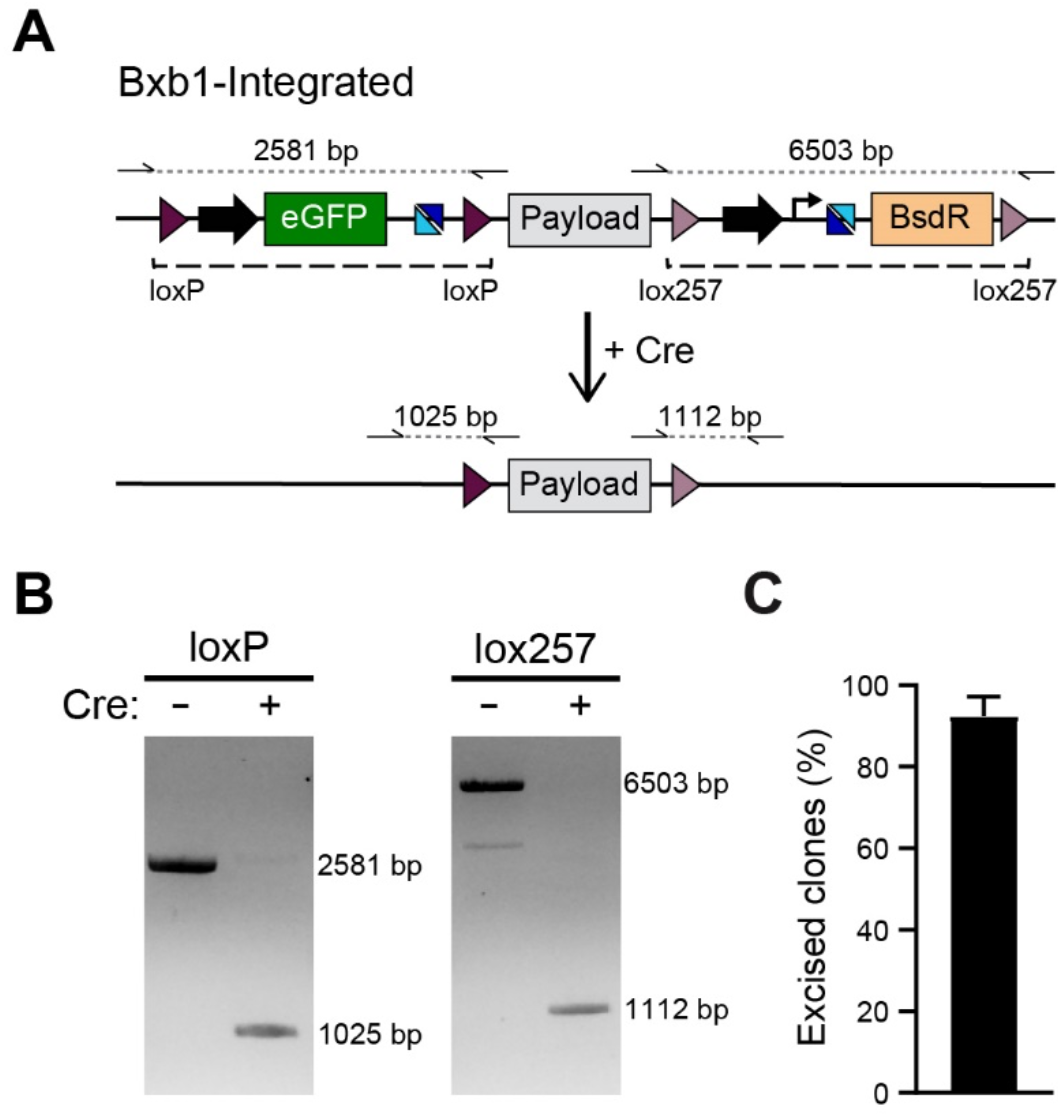
Excision of auxiliary sequences by Cre recombinase. (A) Schematic of procedure for excising the positive selection cassettes and vector backbone following integration of the donor vector into the Bxb1-LP. Dashed lines indicate the sequences excised. Half arrows indicate primer binding sites with dotted lines representing the resulting PCR amplicons. (B) PCR screening using primer pairs indicated in (A), confirming the reduction in amplicon length upon expression of Cre recombinase (+). (C) Quantification of integrated hiPSCs that have excised the auxiliary sequences following Cre expression. n=5 independent transfections; error bars represent ± SEM.

Overall, combining Bxb1-mediated integration of DNA vectors with subsequent Cre-mediated excision of the auxiliary sequences resulted in targeted clonal hiPSC lines being generated within 6 weeks, irrespective of the size of the DNA integrated. Furthermore, due to the efficiency of the recombinases combined with drug selection, correctly targeted cells were identified when screening typically <10 clones.

### STRAIGHT-IN expedites evaluating and generating multi-parameter reporter hiPSCs

To demonstrate the utility and rapid adaptability of STRAIGHT-IN, we generated a series of hiPSC lines to initially evaluate individual optogenetic sensors prior to developing a multi-parameter reporter line for assessing excitation-contraction coupling in hiPSC-CMs. These reporters consisted of ASAP2f for assessing the cardiac action potential (AP) (Zhang et al., 2019), jRCaMP1b for measuring cytosolic Ca^2+^ levels (Dana et al., 2016), and a far-red fluorescent reporter fused to a plasma membrane-localisation signal (Lck-miRFP703) (Chertkova et al., 2017; Shcherbakova et al., 2016) to quantify contraction. To simplify construction of the donor vectors, a modular cloning strategy was employed, in which the basic components (e.g. promoter, localisation signal, coding DNA sequence (CDS), terminator) were initially constructed in a one-step golden gate cloning reaction (Weber et al., 2011). The resulting expression cassettes could then be assembled either individually or as a complex multi-unit circuit into a modified donor vector.

Each reporter was individually integrated into the *AAVS1-*Bxb1 hiPSCs and auxiliary sequences excised, with targeted clones confirmed by genotyping PCR (**Figures S4A and S4B**). Fluorescence imaging established that the reporters were constitutively expressed either in the cell membrane (ASAP2f, miRFP703) or the cytosol (jRCaMP1b) of hiPSCs (**Figure S4C**). All three hiPSC lines differentiated into hiPSC-CMs which displayed characteristic sarcomeric structures, as evidenced by α-actinin staining (**Figure S4D**), and the reporters remained constitutively expressed and localised to the expected subcellular regions (**Figure S4E**). Furthermore, these fluorescent sensors facilitated the assessment of APs, Ca^2+^ transients and contraction in hiPSC-CMs (**Figure S4F**). The hiPSC-CMs expressing ASAP2f showed the expected periodic changes in fluorescence intensity, with a reduction detected in the depolarization phase, followed by an increase during membrane repolarization and the diastolic resting phase. Likewise, jRCaMP1b-expressing hiPSC-CMs displayed cyclic changes in fluorescence, with an increase in fluorescence intensity during the systolic rise in intracellular calcium levels, followed by a reduction in fluorescence during relaxation. Finally, contraction could be quantified by measuring the displacement of membrane-localised miRFP703 (Sala et al., 2018). Accordingly, we constructed a donor vector comprising all three reporters plus an additional copy of ASAP2f to increase its expression in hiPSC-CMs (**Figure S4G**). Using STRAIGHT-IN, we integrated the multi-reporter construct into the *AAVS1-*Bxb1 hiPSCs (∼35% efficiency) and excised the LP cassette and vector backbone (**Figures S4H and S4I**). The resulting hiPSC line (*AAVS1*-AJMA), containing an ∼14 kb DNA payload, co-expressed all 3 reporters in >90% of the cells (**Figures 4A and 4B**). All 3 reporters were also expressed in hiPSC-CMs, although with some silencing of jRCaMP1b and Lck-miRFP703 (**Figures 4C-E**). Nevertheless, excitation-contraction coupling could be assessed using these sensors (**Figure 4F**).

**Figure 4.**
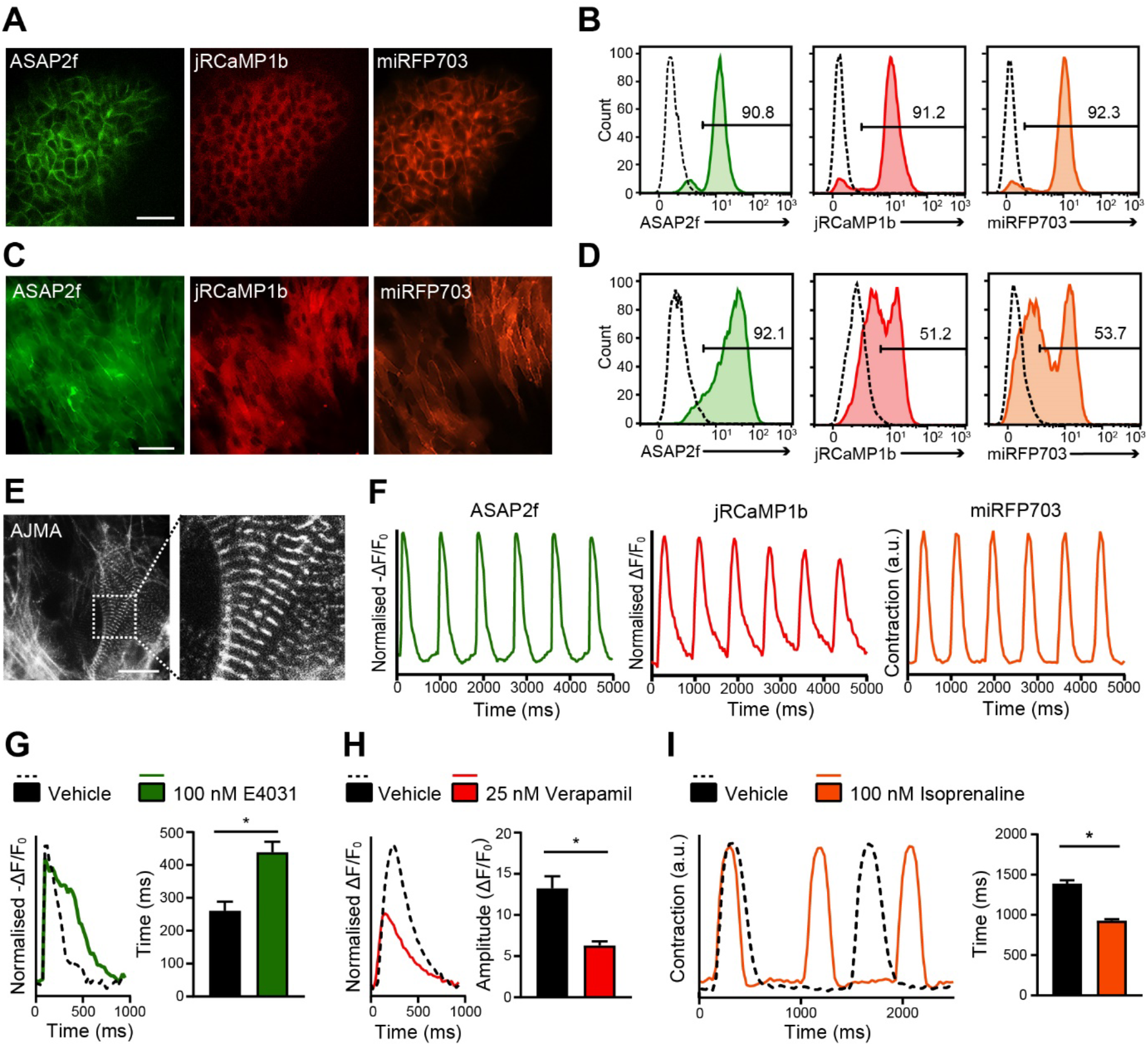
Generation of a multi-parameter reporter hiPSC line using STRAIGHT-IN. (A – D) Fluorescence images (A, C) and flow cytometric analysis (B, D) of *AAVS1*-AJMA hiPSCs (A, B) and hiPSC-CMs (C, D) indicating the cellular localisation and expression level of each of the reporters integrated. Scale bars, 75 µm. Numbers in histograms indicate the percentage of cells positive for the specified reporter; dotted lines indicate negative control. (E) Immunofluorescence image of the cardiac sarcomeric protein α-actinin in *AAVS1-AJMA* hiPSC-CMs. Image on the right is a magnification of the region within the dotted boxes: Scale bar, 25 μm. (F) Representative time plots of baseline-normalised fluorescence signals from *AAVS1*-AJMA hiPSC-CMs stimulated at 1.2 Hz. Changes in the fluorescence of ASAP2f (*left*) and jRCaMP1b (*middle*) reflect the action potential and cytosolic Ca^2+^ transients respectively, while displacement of the miRFP703 (*right*) fluorescence signal indicates contraction dynamics. (G-I) Representative AP (G, *left*), cytosolic Ca^2+^ (H, *left*) and contraction (I, *left*) transients of *AAVS1*-AJMA hiPSC-CMs treated with vehicle (0.1% DMSO) or indicated compounds, together with the resulting average APD_90_ (G, *right*), Ca^2+^ peak amplitude (H, *right*) contraction peak-to-peak time (I, *right*) values. n=4 (*vehicle*) and n=5 (*drug*) treated samples; error bars represent ± SEM; * p < 0.01 (unpaired t-test).

Furthermore, we evaluated the ability of *AAVS1*-AJMA cardiomyocytes to detect changes in AP, intracellular Ca^2+^ transients and contraction profiles for cardiac safety pharmacology applications. For electrophysiological responses, the hiPSC-CMs were treated with the specific hERG channel blocker, E-4031. Compared to the vehicle control, significant prolongation in AP duration at 90% repolarisation (APD_90_) was observed following addition of E-4031 (**Figure 4G**). Similarly, changes in Ca^2+^ handling were detected in the presence of verapamil, a multi-channel blocking compound. Consistent with its mechanism as an L-type Ca^2+^ channel blocker, a decrease in Ca^2+^ transient amplitude was observed (**Figure 4H**). Finally, addition of the beta-adrenergic agonist isoprenaline resulted in a shortening of the contraction duration of the hiPSC-CMs and a significant decrease in mean cycle length (**Figure 4I**).

Together, these results demonstrate how STRAIGHT-IN, combined with adaptations to the donor vector for modular cloning and the development of a component library, can be applied to rapidly construct and integrate synthetic genetic circuits, thus permiting the comparison of various transgenes under the same chromosomal environment to exclude position effects. The stable integration of these reporters into a genomic safe-harbor locus minimises the unpredicted consequences of viral-based approaches and offers the possibility to repeatedly measure the hiPSC-CMs over multiple timepoints, for example to monitor the maturation of the cells or to evaluate chronic pharmacological responses (Karbassi et al., 2020; De Korte et al., 2020).

### STRAIGHT-IN facilitates the simultaneous generation of a panel of disease variant hiPSCs

Lastly, we investigated whether the platform supported multiplex genetic assays by simultaneously generating a library of hiPSC lines carrying heterozygous mutations in *KCNH2*. Mutations in *KCNH2*, which encodes the hERG ion channel, can cause various cardiac arrhythmia syndromes including long QT syndrome type 2 (LQT2), short QT type 1, and Brugada syndrome (Chen et al., 2016). However, there are also rare *KCNH2* variants that do not affect the functionality of the encoded ion channel (Ng et al., 2020). Determing whether these variants are disease-causing or innocuous is critical for determining the best course of action for treating the patient (Giudicessi et al., 2018).

As there appears to be no limit on the length of the DNA sequence that can be inserted using STRAIGHT-IN, we opted to replace the entire *KCNH2* genomic locus (∼54 kb) on one allele with the Bxb1-LP (**Figure 5A**). This meant that the introduced heterozygous *KCNH2* variants would be in an almost identical genomic context to that in affected individuals. Genotyping and quantitative PCRs confirmed that the resulting hiPSC line (*KCNH2*^+/Acc^) was correctly targeted, contained a single integration of the Bxb1-LP and was monoallelic for *KCNH2* while retaining two copies of the flanking genes, *NOS3* and *AOC1* (**Figures 5B and 5C**).

**Figure 5.**
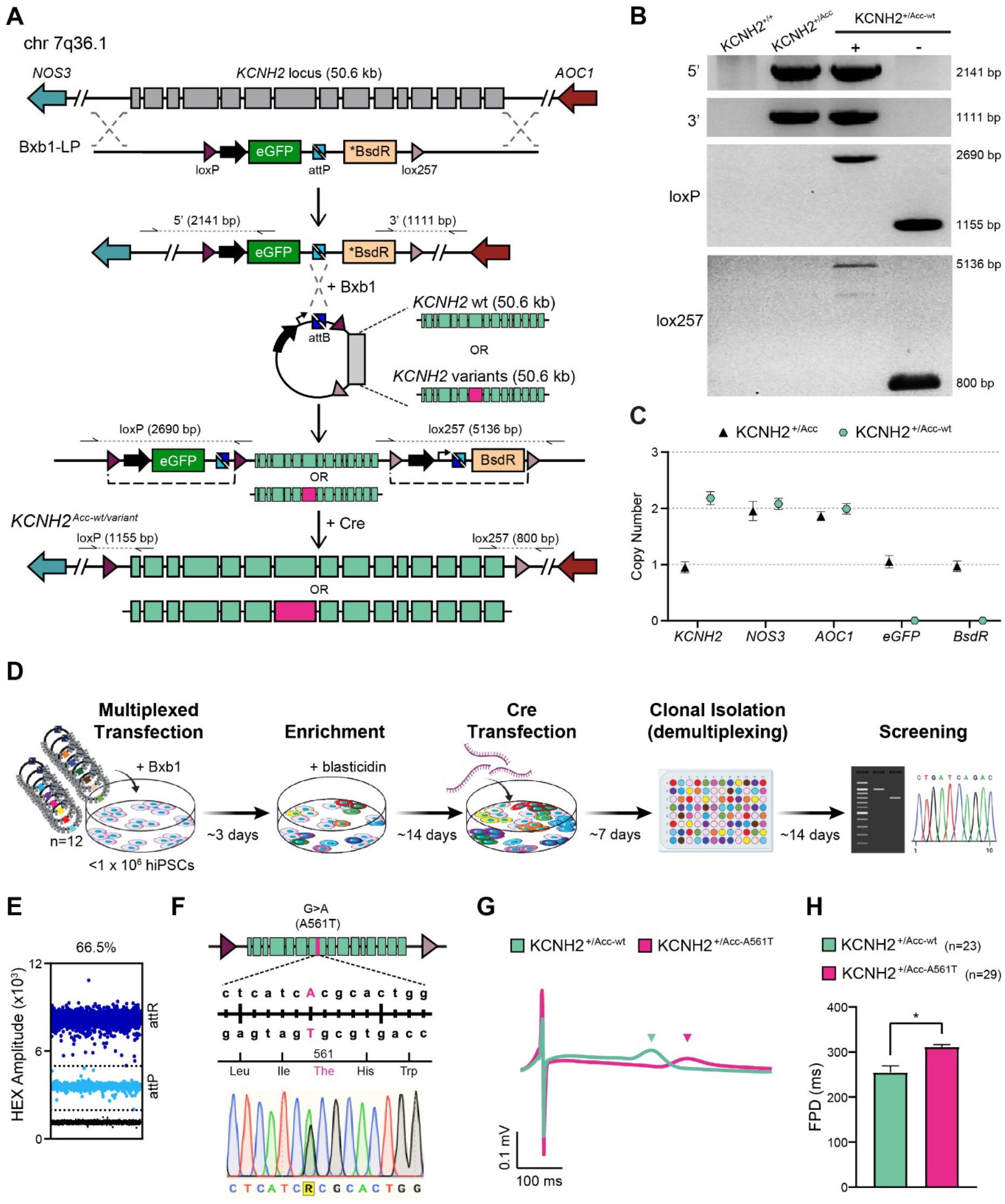
Simultaneously generating a panel of *KCNH2*-variant hiPSC lines with STRAIGHT-IN. (A) Schematic of STRAIGHT-IN procedure to perform targeted heterozygous modifications to a 50.6 kb genomic region on chromosome 7 that includes *KCNH2*. Half arrows indicate primer binding sites with dotted lines representing the resulting PCR-generated amplicons. Dashed lines indicate the sequences excised by Cre recombinase. (B) PCR products amplified with the corresponding primer pairs indicated in (A), confirming targeting of Bxb1-LP to *KCNH2* (*KCNH2*^+/Acc^), and subsequent re-introduction of wildtype *KCNH2* (*KCNH2*^+/Acc-wt^). Auxiliary sequences detected upon integration (+) were excised following Cre expression (-). The sizes of the amplicons are indicated. (C) ddPCR confirming that *KCNH2*^+/Acc^ and *KCNH2*^+/Acc-wt^ hiPSCs contained the expected number of copies of the genomic genes *KCNH2* (1 and 2 copies respectively), *NOS3* and *AOC1* (both 2 copies), and the Bxb1-LP cassette transgenes, *eGFP* and *BsdR* (1 and 0 copies respectively). Error bars represent Poisson 95% CI. (D) Schematic of the STRAIGHT-IN procedure for simultaneously generating and identifying isogenic hiPSC clones for 12 different *KCNH2* variants, along with the approximate time required for each step. (E) Dot plot of *KCNH2*^*+/Acc*^ hiPSCs transfected with the 12 *KCNH2* variant donor vectors. Dots represent droplets containing the indicated sequence (attR or attP), while the percentage denotes the calculated integration efficiency. (F) Overview of the genomic sequence and location within *KCNH2* of the introduced variant A561T, and sequence analysis from one of the resulting *KCNH2*^*+/Acc-A561T*^ hiPSC clones indicating the heterozygous introduction of c.G1681A. (G,H) Representative averaged field potential (FP) traces (G) and averaged FP duration (FPD) values (H) of *KCNH2*^+/Acc-wt^ and *KCNH2*^+/Acc-A561T^ hiPSC-CMs paced at 1.25 Hz. Colored arrowheads indicate the respective repolarization peak for each line. Values (n) indicate the number of recordings. Error bars represent ± SEM, and * p < 0.0001 (unpaired t-test).

We initially reintroduced the *KCNH2* wild type sequence into the *KCNH2*^+/Acc^ hiPSC line as confirmed by PCR screening (**Figure 5B**). From 59 clones screened, 18 (∼30%) had undergone integration and excision, showing similar efficiency to that observed for the *AAVS1* locus. Quantitative PCR confirmed for one of the hiPSC clones (*KCNH2*^+/Acc-wt^) that it was now biallelic for *KCNH2*, while both *eGFP* and *BsdR* were absent (**Figure 5C**). Furthermore, *KCNH2*^+/Acc-wt^ was karyotypically normal and differentiated into hiPSC-CMs with similar efficiency as the original wildtype hiPSCs (KCNH2^+/+^) (**Figures S5A-C**). Additionally, no differences in cardiac field potential duration (FPD) were observed between the two lines, indicating that the electrophysiological activity of the hiPSC-CMs was also unaffected by the STRAIGHT-IN procedure (**Figures S5D and S5E**).

Based on this, we constructed donor vectors for 12 *KCNH2* variants identified in exon 7 on ClinVar (**Table S1**), that were subsequently pooled and transfected along with Bxb1 into <1×10^6^ KCNH2^+/Acc^ hiPSCs (**Figure 5D**). Modifications to the enrichment step improved the proportion of recombined hiPSCs to 66.5% (**Figure 5E**), with all 12 variants detected within this mixed population (**Figure S5F**). Following Cre-mediated excision of the auxiliary sequences, 11 out of the 12 variants were recovered from screening 208 colonies (**Figure S5F**), with the entire procedure taking ∼2 months.

We further characterized an hiPSC clone identified as being heterozygous for the *KCNH2* variant A561T (*KCNH2*^+/Acc-A561T^) (**Figure 5F**). Quantitative PCR verified STRAIGHT-IN had occurred as expected and the hiPSCs had a normal karyotype (**Figures S5G and S5H**). hiPSC-CMs that carry this mutation exhibit a prolonged FPD (Brandão et al., 2020; Matsa et al., 2011), reflecting the electrophysiological characteristics of LQT2. The *KCNH2*^+/Acc-A561T^ hiPSCs were differentiated with the resulting CMs having a significantly prolonged FPD compared to *KCNH2*^+/Acc-wt^ hiPSC-CMs (**Figures 5G and 5H**), thereby confirming that *KCNH2* variant models generated using the STRAIGHT-IN procedure likewise can exhibit the expected disease phenotype.

Overall, these results demonstrate how STRAIGHT-IN also can be used as a high-throughput method to multiplex and simultaneously insert potentially hundreds of different disease-linked variants into a control hiPSC line.

## DISCUSSION

In this study we present STRAIGHT-IN, an efficient and modular platform for targeted genomic integration of DNA payloads into hiPSCs. The workflow of STRAIGHT-IN consists of three steps: (i) targeting a LP cassette to the locus of interest; (ii) integrating a donor vector encoding the DNA payload into the LP cassette via a serine recombinase, and; (iii) expressing a tyrosine recombinase to excise the majority of the accessory exogenous DNA sequences. The resulting hiPSC lines contain the targeted DNA payload, with minimal traces of unrequired sequences remaining in the modified locus.

With STRAIGHT-IN we successfully targeted two separate loci in hiPSCs with similar efficiencies, indicating the procedure is not locus dependent and could likely be used to modify any locus of interest. Additionally, two distinct LP cassettes and donor constructs were designed that utilised different serine and tyrosine recombinases. Although we have focussed on the capabilities of Bxb1-mediated recombination, the specificity of the recombinases also enables dual modification of a cell line using STRAIGHT-IN, thereby further broadening the flexibility of the system. Additional modifications to the LP cassettes, such as the inclusion of mutant *attP* and *attB* sites that result in better integration efficiencies (Jusiak et al., 2019), would likely improve the efficiency of the procedure even more.

Our results also suggest that there is no constraint on the size of the DNA payload that can be integrated, overcoming one of the main limitations of other commonly-used DNA integration systems (e.g. viral vectors, programmable nuclease knock-in). To date, the largest payloads reported to be integrated into a single artificial LP in mammalian cells was ∼33 kb in CHO cells and <10 kb in hiPSCs (Gaidukov et al., 2018; Zhu et al., 2014). Here, we were readily able to integrate DNA sequences varying between ∼14 – 50 kb in length, and even managed to integrate a donor vector containing a 170 kb+ BAC fragment. Subsequent tyrosine recombinase-mediated excision of the LP cassette and donor vector backbone was also very efficient with >90% of the integrated cells having these auxiliary sequences excised. This step of the procedure also could be performed by transfecting the recombinase as mRNA, ensuring expression was transient and eliminating the risk of insertional mutagenesis.

For various research and clinical applications, there is an increasing need to precisely integrate large DNA fragments (Zhang et al., 2021). STRAIGHT-IN could potentially simplify the generation of cell lines or animal models containing these large and complex genetic circuits. Here we demonstrate using STRAIGHT-IN to rapidly prototype synthetic genetic circuits and as a high-throughput method to simultaneously generate a panel of hiPSC lines for modelling a genetic disease. Moreover, this platform could also be used as a tool for gene therapy and in the assembly of custom-designed mammalian chromosomes (Hessel et al., 2016).

We envision that STRAIGHT-IN will be especially valuable for the large-scale generation of disease panels for precision medicine applications. Such panels could be used to test the efficacy of pharmacological compounds against individual mutations. For example, nearly 500 mutations in *KCNH2* have been associated with LQT2 with at least 170 of these predicted to cause trafficking defects (Anderson et al., 2014). Orkambi, an approved drug for treating cystic fibrosis in certain patients, was identified as a potential therapeutic for patients with LQT2 caused by trafficking defective variants and is now being clinically evaluated (Schwartz et al., 2019; Wu et al., 2020). However, for some variants lumacaftor, one of the compounds present in Orkambi, appeared to cause an opposite effect (O’Hare et al., 2020). Therefore, large scale efficacy studies of the drug using a cohort of *KCNH2* variant hiPSC-CMs will likely be required to advance this further in the clinic.

Another purpose that would benefit from this multiplex approach is classifying rare variants. Large scale genetic sequencing projects have revealed that rare variants are highly prevalent in the general population. The difficulty in distinguishing pathogenic variants from rare benign variants when performing genetic testing for inherited disorders has resulted in large proportions of patients having variants classified as being of “uncertain significance” and so not clinically actionable. Therefore, platforms that are rapid and do not require an individualized targeting strategy for each variant would be highly advantageous in a diagnostic setting. A previous study investigated whether a dual integrase cassette exchange strategy could generate such a hiPSC panel of *TNNT2* variants (Lv et al., 2018). While heterozygous clones for 14 coding variants were isolated, the reported efficiency was ∼5%, and these lines represented only 12% of the variants introduced. Additionally, the procedure was restricted to integrating a DNA payload of < 1 kb and so only a partial cDNA spanning *TNNT2* exons 6 to 17 could be examined. Not only did this prevent all *TNNT2* CDS variants being investigated, but potentially also resulted in the loss of regulatory elements that controlled expression of the variant. Although we modified a different genetic locus, we reported a much higher targeting efficiency (∼34%) with STRAIGHT-IN and could recover >90% of the variants introduced. Furthermore, with our platform, non-coding genomic regions are retained in the integration. This also permits SNPs identified in genome-wide association studies that might influence the disease phenotype to be modified and investigated.

## Limitations of the study

STRAIGHT-IN is most suitable when the same genomic region will be repeatedly modified or when the DNA cargo for targeting exceeds 5 kb. Other targeting approaches, such as those mediated by programmable nucleases, are likely to be more appropriate if only a few independent modifications (e.g. <4) to a single genomic locus are anticipated, or if the disease variants being investigated are in separate, distinct loci.

To date we have only established the platform in human iPSCs, however we believe this approach is broadly applicable for use in other cell types such as adult stem cells (Menche and Farin, 2021). In some instances, the LP cassette, which can be easily customised, might require modifying to provide alternative approaches for isolating the integrated clones (for example via cell surface markers). Likewise, the current method for excising the unrequired sequences flanking the DNA payload following integration results in traces of these sequences remaining in the modified locus (<300 bp). These remaining sequences are beneficial as they simplify screening procedures, and currently the resulting cell lines are assessed to confirm that the residual auxiliary sequences do not affect expression from the modified locus. However recent methods developed for scarless excision potentially could enable the complete removal of these sequences (Li et al., 2013; Roberts et al., 2019).

## METHODS

Unless otherwise stated, cell culture reagents were obtained from ThermoFisher, while oligonucleotides, hydrolysis probes and synthesised DNA fragments (gBlocks and eBlocks) were obtained from Integrated DNA Technologies (IDT).

### hiPSC line culture

The hiPSC line LUMC0020iCTRL-06 (female, (Zhang et al., 2014)) was generated from primary skin fibroblasts using Sendai virus by the LUMC hiPSC core facility. This line and the resulting subclones used in downstream experiments were assessed for pluripotency, routinely tested for mycoplasma and karyotyped by G-banding. Chromosome spreads and analyses were performed by the Laboratory for Diagnostic Genome Analysis (Leiden University Medical Center). For each cell line, 20 metaphase spreads were examined with samples of sufficient quality to detect numerical and large structural abnormalities. For alkaline phosphatase staining, the alkaline phosphatase detection kit (Merck) was used following manufacturer’s instructions.

All hiPSC lines were maintained in StemFlex™ Medium on laminin-521 (LN521; BioLamina)-coated (1.5 μg/cm^2^) plates. Cells were passaged twice a week by dissociating with either 1x TrypLE Select or Accutase® solution (Sigma).

### hiPSC transfections

Intracellular delivery of DNA, RNA or protein into hiPSCs was accomplished by either electroporation or lipofection using conditions previously described (Brandão et al., 2021). Electroporation was used to deliver Cas9-gRNA RNP complexes along with targeting constructs, or for delivering the BAC_attB(bxb) vector. All other transfections were performed using Lipofectamine™ Stem Transfection Reagent.

### hiPSC subcloning

Dissociated hiPSCs were filtered to remove cell aggregates before being clonally isolated using the single-cell deposition function of a BD FACSAria™ III (BD Biosciences). Here, single hiPSCs were deposited directly into each well of an LN521-coated (1.8 μg/cm^2^) 96-well plate. Media was changed every 3 days for 2 weeks, after which the cells were replicated for screening and archiving (Brandão et al., 2021).

### Genomic DNA (gDNA) extraction

For hiPSCs cultured in 96 well-plates, gDNA was extracted using QuickExtract™ (Lucigen). Cells were resuspended in 30 μl QuickExtract and incubated at 65°C for 15 min, followed by 68°C for 15 min and 98°C for 10 min. For hiPSCs cultured in other formats, gDNA was extracted using the High Pure PCR Template Preparation Kit (Roche) according to the manufacturer’s instructions.

### Cas9 RNP & sgRNA synthesis

Cas9 protein was either purchased (IDT) or kindly provided by N. Geijsen (D’Astolfo et al., 2015). Candidate gRNAs with high specificity were identified around the intended mutation site using the bioinformatics tool, CRISPOR (Concordet and Haeussler, 2018), or were previously published (Wang et al., 2015). The gRNAs were synthesised as chimeric single gRNAs (sgRNAs) by in vitro transcription.

### Bxb1-LP and φC31-LP cassette construction

The components of both cassettes were first PCR-amplified from other vectors and subsequently ligated together using the NEBuilder® HiFi DNA Assembly Cloning kit (NEB) to generate the resulting pENTR-eGFP-attP(bxb)-*BsdR and pENTR-mCherry-attP(C31)-*BleoR vectors. Briefly, pENTR-eGFP-attP(bxb)-*BsdR was composed of the backbone of the cloning vector pENTR/D-TOPO, a *loxP* sequence together with a PGK promoter, an *eGFP* reporter, and a blasticidin resistance gene lacking an initiation codon (\**BsdR*). Primers were used to introduce the *lox257* and Bxb1-specific *attP* sequences into the final vector. Similarly, pENTR-mCherry-attP(C31)-*BleoR consisted of the same backbone vector, an *FRT* sequence together with a PGK promoter, a *mCherry* reporter, and a bleomycin resistance gene lacking an initiation codon (\**BleoR*). Again, primers were used to introduce the *F3* and φC31-specific *attP* sequences into the final vector.

### *AAVS1-*Bxb1, *AAVS1-*φC31 and *AAVS1-*Dual hiPSC line generation

To target the Bxb1-LP and φC31-LP cassettes to the adeno-associated virus integration site (*AAVS1*) within intron 1 of *PPP1R12C*, the cassettes were cloned via NEBuilder® HiFi DNA Assembly into AAVS1_SA_2A_Neo_CAG_RTTA3 (Addgene, #60431) (Sim et al., 2014). The resulting targeting vectors (AAVS1-Bxb1-LP-TC and AAVS1-φC31-LP-TC) therefore had the Bxb1-LP and φC31-LP cassettes flanked by ∼800 bp homology arms. Either AAVS1-Bxb1-LP-TC and/or AAVS1-φC31-LP-TC along with Cas9-*AAVS1* gRNA RNP complex (gRNA: 5’-GGGGCCACTAGGGACAGGAT-3’) were electroporated into LUMC0020iCTRL-06 hiPSCs. Following recovery and expansion of the electroporated cells, eGFP^+^, mCherry^+^ or double-positive hiPSCs were clonally isolated. Targeted clones were identified by PCR screening over the 5’ and 3’ homology arms.

### Bxb1 and φC31 donor cloning vector construction

Three Bxb1 donor vectors were constructed for the cloning of various DNA payloads. For inserting DNA payloads <20 kb, the pBR_attB(bxb)_lox cloning vector was used (**Figure S1C, *left***). The cloning vector, p15_attB(bxb)_lox, was constructed for the cloning of 20 – 50 kb DNA payloads (**Figure S1C, *middle***). Finally, the cloning vector, pBR_attB(bxb)_ccdB_lox, was developed for modular construction of multi-component synthetic circuits (**Figure S1C, *right***).

The φC31 donor vector, pBR_attB(C31)_FRT, was constructed by digesting pEFBOS_creIRESBsd with EcoRI (NEB) and ligating with a gBlock that contained a φC31-specific *attB* site, as well as *FRT* and *F3* sequences (**Figure S1D**).

### Vectors for evaluating the integration of different DNA payload sizes

To insert DNA payloads between ∼2 – 50 kb into either the pBR_attB(bxb)_lox or p15_attB(bxb)_lox donor vectors, fragments from a BAC carrying the human *KCNH2* gene (RP11-10L20) were subcloned via recombineering (**Figure S1E**) (Fu et al., 2010).

To modify RP11-10L20 to enable targeted integration of the complete BAC construct (Bxb1-BAC donor) into the *AAVS1-*Bxb1 hiPSCs, recombineering was used to insert a DNA fragment that included the EF1a promoter, Bxb1-*attB* and a kanamycin-resistance cassette.

### Integrating donor vectors into the *AAVS1* acceptor hiPSC lines

Unless stated otherwise, 1.2 μg of the donor vectors along with 0.8 μg of the corresponding integrase-expressing plasmids, pCAG-NLS-HA-Bxb1 (Addgene #51271) (Hermann et al., 2014) and pCAG-φC31 (Addgene #62658) (Ohtsuka et al., 2015), were transfected by lipofection into *AAVS1-*Bxb1, *AAVS1-*φC31 and *AAVS1-*Dual hiPSC lines. To integrate the modified RP11-10L20 BAC construct into the *AAVS1-*Bxb1 hiPSCs, Bxb1-BAC donor was co-electroporated with pCAG-NLS-HA-Bxb1. For both approaches, ∼3 days after transfection the cells were harvested and passaged. To enrich for integrated hiPSCs, either blasticidin (2 μg/ml, Sigma) or zeocin (15 μg/ml, ThermoFisher) were added to the culture medium for a period of 12 and 5 days, respectively. Donor vector integration was confirmed via a PCR screening strategy to detect the formation of the two new recombination sites, *attR* and *attL*.

### Excising the auxiliary sequences

hiPSCs were transfected by lipofection with either an Flp-expressing plasmid (1.6 μg, pCAG_FlpoIRESpuro (Kranz et al., 2010)), a Cre-expressing plasmid (1.6 μg, pEFBOS_CreIRESpuro (Davis et al., 2008)) or StemMACS™ Cre Recombinase mRNA (200 ng, Miltenyi Biotec). For hiPSCs transfected with the plasmids, selection with puromycin (1 μg/ml, Sigma) was initiated 24 h post-transfection and maintained for 48 h. Genotyping PCR was used to confirm that the *loxP*- and *lox257*- or the *FRT*- and *F3*-flanked sequences were excised.

### Droplet digital PCR (ddPCR)

ddPCR was performed and analysed using a thermocycler, the Q200 AutoDG and QX200™ Droplet Digital PCR System, and QuantaSoft software (all Bio-Rad). Assays comprising of premixtures of a forward and reverse primer (18 μM each) with a FAM- or HEX-conjugated hydrolysis probe (5 μM) were either purchased from Bio-Rad, based on previous publications (Roberts et al., 2017), or designed based on pre-defined criteria (Bell et al., 2018). Reactions (22 μl) were prepared with 2x Supermix for probes with no dUTP (Bio-Rad), 900 nM of each primer, 250 nM of each probe, and 30-100 ng of gDNA digested with 2-5 U of HindIII, HaeIII or MseI (all NEB) depending on the sequence of the amplicon. Droplet generation, PCR amplification and analysis were all performed according to the manufacturer’s instructions. For CNV assays, the two-copy autosomal gene *RPP30* gene was used as a reference.

*attP:attR assay*. To determine the recombination efficiency of the donor plasmids into the Bxb1-LP or φC31-LP, probes were designed to detect the *attP* or *attR* sites in the transfected hiPSCs. To optimise amplification conditions and confirm the specificity of the assay, gBlocks matching the two expected amplicons for each integrase were mixed in differing ratios and used as template DNA, with a strong correlation (R^2^= >0.99) seen between the expected and observed frequencies (Figure S2).

*KCNH2 variants*. Probes were designed for each of the 12 *KCNH2* variants. To improve target specificity, 2–5 locked nucleic acids were included per probe. Amplification conditions for each probe were optimised for discriminating amplicons containing that specific missense mutation. Droplets were analysed using the “absolute quantification” option of the QuantaSoft software.

### Generating the optogenetic reporter hiPSC lines

The assembly of the individual and multi-parameter reporter donor vectors was based on the modular and hierarchal cloning system, MoClo (Weber et al., 2011). Briefly, the required components for the expression of these reporters (i.e. promoter sequence, localisation signal, CDS and polyA signal) were PCR-amplified using primers that incorporated the recognition site for the type IIS enzyme BsaI and previously designated overhang sequences for positioning and orientation of the components (Andreou and Nakayama, 2018; Weber et al., 2011).

The components for each of the individual optogenetic sensors were first assembled in Level 1 destination vectors included in the MoClo Toolkit (Addgene #1000000044) by Golden Gate (GG) assembly with the restriction enzyme, FastDigest Eco31I (ThermoFisher), thereby generating the intermediate Transcriptional Unit (TU) vectors, which were transformed into 10-beta Competent *E. coli* (NEB). The reporter construct for ASAP2f was assembled into 2 different Level 1 destination vectors, while the constructs for jRCaMP1b and miRFP703 were each assembled into Level 1 destination vectors for positions 2 and 3 respectively.

The resulting TU vectors were subsequently assembled either individually or in combination into the donor vector pBR_attB(bxb)_ccdB_lox by digestion with BpiI. Dummy and end-linker vectors from the MoClo Toolkit were included as required in the GG assembly reaction. The ensuing donor vectors (Bxb1-ASAP2f; Bxb1-jRCaMP1b; Bxb1-miRFP703 and Bxb1-AJMA) were transformed into Stbl2™ *E. coli* for ccdB counterselection.

Each donor vector was separately integrated into the *AAVS1-*Bxb1 hiPSCs, followed by excision of the auxiliary sequences. Clonal hiPSC lines were derived for the single reporter targeted cells, while enrichment for the *AAVS1*-AJMA hiPSCs was performed by flow cytometric sorting of cells co-expressing the 3 reporters. Genotyping PCRs confirmed targeted integration, as well as no rearrangement within the TUs of the *AAVS1*-AJMA hiPSCs.

### Generating the *KCNH2*^+/Acc^ hiPSC line

The vector to target the Bxb1-LP cassette to the *KCNH2* locus (*KCNH2*-Bxb1-LP-TC) was generated by PCR-amplifying the Bxb1-LP cassette with primers containing 80 bp overhangs complementary to endogenous sequences ∼8.5 kb (5’) and ∼8.7 kb (3’) of *KCNH2*. The resulting PCR product was cloned into pMini T2.0 (NEB).

One copy of *KCNH2* was deleted by electroporating Cas9 protein together with two gRNAs targeting both ends (gRNA for 5’ end: 5’-ATGAAGGCTTTCCCATCCGT-3’ and gRNA for 3’ end: 5’-ACTGTGCTGGGTACGCTGAC-3’) into LUMC0020iCTRL-06. Next, the *KCNH2*-Bxb1-LP-TC along with a Cas9-*KCNH2* gRNA RNP complex (gRNA: 5’-CTGGTTGTGCTGACTGTGCT-3’) was electroporated into this modified hiPSC line. Following recovery and expansion of the electroporated cells, eGFP^+^ hiPSCs were clonally isolated. Targeted clones (*KCNH2*^+/Acc^) were further characterised by PCR screening and Sanger sequencing over the 5’ and 3’ homology arms. The resulting *KCNH2*^+/Acc^ hiPSC line selected contained a single integration event of the Bxb1-LP cassette as determined by ddPCR.

### Constructing *KCNH2* wildtype and variant donor vectors

The donor vectors containing the various *KCNH2* genomic sequences were built based on recombineering strategies previously described (Fu et al., 2010; Wang et al., 2014). Briefly, to seamlessly introduce the variants into *KCNH2*, a counterselection cassette (ccdB-Amp) was first introduced to replace exon 7 in the BAC, RP11-10L20. Next, synthetic double-stranded DNA fragments that introduced specific missense mutations in exon 7 of *KCNH2* (Table S1) were pooled and amplified by PCR, before being electroporated into the bacteria to replace the counterselection cassette. Colonies that subsequently grew in the absence of L-arabinose were then screened by PCR and Sanger sequencing to identify recombined BACs for each of the 12 mutations.

The sequence in the BAC corresponding to the *KCNH2* genomic region deleted in the *KCNH2*^+/Acc^ hiPSCs was subsequently subcloned from both wildtype RP11-10L20 as well as clones carrying the introduced variants into p15_attB(bxb)_lox by recombineering. The resulting colonies were screened by PCR to confirm subcloning and the plasmids retransformed to ensure the resulting *KCNH2* donor vectors were pure. Sanger sequencing also confirmed the presence of each missense mutation, and that the *attB* and *lox* sequences were correct. Finally, the integrity of each of the donor vectors was evaluated by restriction analysis.

Bacterial cultures of the *KCNH2* variant donor vectors were pooled and grown at 30°C, while the KCNH2 wildtype donor vector (p15-attB_*KCNH2*_wt_donor) was cultured separately. Plasmid DNA was purified using the NucleoBond® Xtra Midi Kit (Macherey Nagel).

### Generating the *KCNH2*^+/Acc-wt^ and *KCNH2*^+/Acc-variants^ hiPSC lines

To generate the *KCNH2*^+/Acc-wt^ line, a similar procedure to that used to generate the *AAVS1*-reporter hiPSCs was performed, except that 3.2 μg of p15-attB_*KCNH2*_wt_donor was transfected into the *KCNH2*^+/Acc^ hiPSCs. Cells were selected with blasticidin for 6 days, followed by transfection of pEFBOS_CreIRESpuro and selection with puromycin. The resulting *KCNH2*^+/Acc-wt^ hiPSCs were clonally isolated by single-cell deposition and identified by genotyping PCR.

To multiplex the generation of the *KCNH2*^+/acc-variant^ lines, 0.6 μg of the pooled 12 *KCNH2* variant donor vectors was transfected. The antibiotic selection strategy was also modified, with the cells maintained in culture medium containing 2 μg/ml blasticidin over 3 passages (14 days). Excision of the auxiliary sequences was performed by Cre recombinase mRNA transfection. Following single-cell deposition, genotyping PCR detected clones that had undergone both integration and excision steps, while Sanger sequencing identified the heterozygous *KCNH2* variant that was introduced.

### Differentiation and culture of hiPSC-CMs

The hiPSCs were differentiated into cardiomyocytes as previously described (Campostrini et al., 2021). One day prior to differentiation (d-1), the hiPSCs were harvested using TrypLE Select and plated onto Matrigel (1:100, Corning)-coated wells in StemFlex™ Medium containing RevitaCell™ Supplement (1:200 dilution). On d0, the cells were refreshed with mBEL medium containing 5 µM CHIR99021 (Axon Medchem). On differentiation d2, the cells were refreshed with mBEL medium containing 5 µM of XAV939 (Tocris) and 0.25 µM IWP-L6 (AbMole). From differentiation d4 on, the cells were maintained in mBEL medium. The hiPSC-CMs were cryopreserved at differentiation d20 or d21 as previously described in a freezing medium comprising of 90% Knockout Serum Replacement (Gibco) and 10% DMSO (Brink et al., 2020). Subsequent thawing and seeding of the cells were performed as previously described (Brink et al., 2020; Campostrini et al., 2021).

### Flow cytometric analysis

A single-cell suspension of hiPSCs or hiPSC-CMs was obtained by dissociating the cells with TrypLE Select and filtering the cell suspension. Cells were fixed and permeabilised using the FIX and PERM™ Cell Permeabilization Kit (Invitrogen) according to manufacturer’s instructions. The hiPSC-CMs were incubated with the conjugated antibodies cTnT-Vioblue or cTnT-FITC (1:50, Miltenyi Biotec, #130-120-402 or #130-119-575). All antibodies were diluted in permeabilization medium (medium B; Invitrogen). The data was acquired using a MacsQuant VYB flow cytometer (Miltenyi Biotec) and analysed using FlowJo software (v. 10.2, FlowJo).

### Fluorescence imaging

The hiPSCs and hiPSC-CMs were seeded on 96-well imaging microplates (Corning), with images of the fluorescent reporters acquired using an EVOS™ M7000 Cell Imaging System (ThermoFisher) at 40x magnification. For visualizing the sarcomeres, the hiPSC-CMs were fixed and permeabilised with the FIX and PERM™ Cell Permeabilization kit and labelled with an antibody specific for α-actinin (1:250, Sigma-Aldrich #A7811), followed by an Alexa Fluor 350-conjugated secondary antibody (1:500, ThermoFisher, #A-11045).

### Optical evaluation of hiPSC-CMs

hiPSC-CMs differentiated from the optogenetic reporter hiPSCs were seeded on 96-well imaging microplates pre-coated with Matrigel (1:100) in mBEL medium. Medium was refreshed the next day and every 2-3 days thereafter, with analysis performed 7 days after thawing. Before performing baseline measurements, wells were refreshed with 200 μL mBEL medium and left for 60 min at 37°C to equilibrate.

After baseline measurements, the hiPSC-CMs were refreshed with 100 μL of mBEL including compounds at final test concentrations and incubated for 5 min at 37 °C before recording. Compounds used were E4031, verapamil (both Tocris Bioscience) and isoprenaline (Sigma-Aldrich). All compounds were reconstituted in DMSO (Sigma-Aldrich), with solutions prepared to ensure a final concentration of 0.1% DMSO in each well. Vehicle incubations were done similarly using mBEL + 0.1% DMSO. Measurements were made with cells paced at 1.2 Hz using a pair of field stimulation electrodes placed in the culture medium, except for the isoprenaline measurements which were performed on spontaneously beating hiPSC-CMs. A Leica DMI6000B imaging system (Leica Microsystems) equipped with 470, 565 and 656 nm lasers was used to record signals for AP, cytosolic Ca^2+^ and contraction transients respectively. The microscope was fitted with an environmental chamber that allowed for the measurements to be performed at 37°C and 5% CO_2_. The analyses for AP and cytosolic Ca^2+^ transients were performed using ImageJ (NIH) and algorithms developed in-house (van Meer et al., 2019). For contraction transients, the analyses were performed using the software, MUSCLEMOTION (Sala et al., 2018).

### Multielectrode array (MEA) recordings of hiPSC-CMs

The hiPSC-CMs were seeded in a Matrigel-coated 96 well Lumos MEA plate (Axion BioSystems, Inc.) in 5 μl mBEL medium supplemented with RevitaCell™ (1:200). The cells were incubated for 1 h at 37°C, 5% CO_2_ to allow attachment, with the wells then supplemented with an additional 150 µl mBEL medium. The medium was refreshed the next day and every 2-3 days thereafter.

Recordings were performed using the Maestro Pro multiwell MEA platform (Axion BioSystems, Inc.). Field potential (FP) recordings were either performed on spontaneously beating hiPSC-CMs (*KCNH2*^*+/+*^ vs *KCNH2*^*+/Acc-wt*^) or on optically paced cells (*KCNH2*^*+/Acc-wt*^ vs *KCNH2*^*+/Acc-A561T*^). Optical pacing of the hiPSC-CMs was performed as previously described (Brink et al., in press). Briefly, at day 8 or 9 post-seeding the hiPSC-CMs were transfected with 50 ng of *in vitro* transcribed Channelrhodopsin-2 (ChR2) mRNA. 2 or 3 days post-transfection, and at least 1 h before recordings, medium was refreshed with mBEL supplemented with 1 μM all-*trans* retinal (Sigma). Subsequently, the hiPSC-CMs were paced at 1.25 Hz using 10 ms pulses of blue light (475 nm) delivered for 5 min using the Lumos™ Optical Stimulation System (Axion Biosystems, Inc.).

Prior to all recordings, the hiPSC-CMs were equilibrated inside the device at 37°C, 5% CO_2_ for 10 min. Recordings were performed for 3-4 min using Cardiac Standard filters and amplifiers in spontaneous cardiac mode (12.5 kHz sampling frequency; 0.1–2000 Hz band pass filter). Raw data files were re-recorded to generate CSV files using the AxIS digital filters. CSV and RAW files were loaded into the Cardiac Analysis Tool (Axion BioSystems, Inc., version 3.1.5) to enable precise assessment and analysis of the FP duration (FPD) from one “golden electrode” per well.

### Statistical analysis

All data are presented as mean, with detailed statistics and statistical significance indicated in the figure legends. The statistical power was determined using the two-tailed t-test for unpaired measurements. Differences were considered statistically significant at *P* <0.05. Sample sizes are indicated in the figures. Statistical analysis was performed with GraphPad Prism 8 software (v8.2.0, GraphPad).

## ACKNOWLEDGEMENTS

We thank M. Bellin for providing the control (LUMC0020iCTRL) hiPSC line, M. de Graaf and L. Voortman (LUMC light microscopy facility) for microscopy assistance and the LUMC flow cytometry facility for sorting the cells. We also acknowledge Francis Stewart for sharing plasmids for recombineering and Niels Geijsen for providing the Cas9 protein. Some panels within figures were created with BioRender.com.

