## Supplemental files for "STRAIGHT-IN: A platform for high-throughput targeting of large DNA payloads into human pluripotent stem cells"

[6]Lead contact

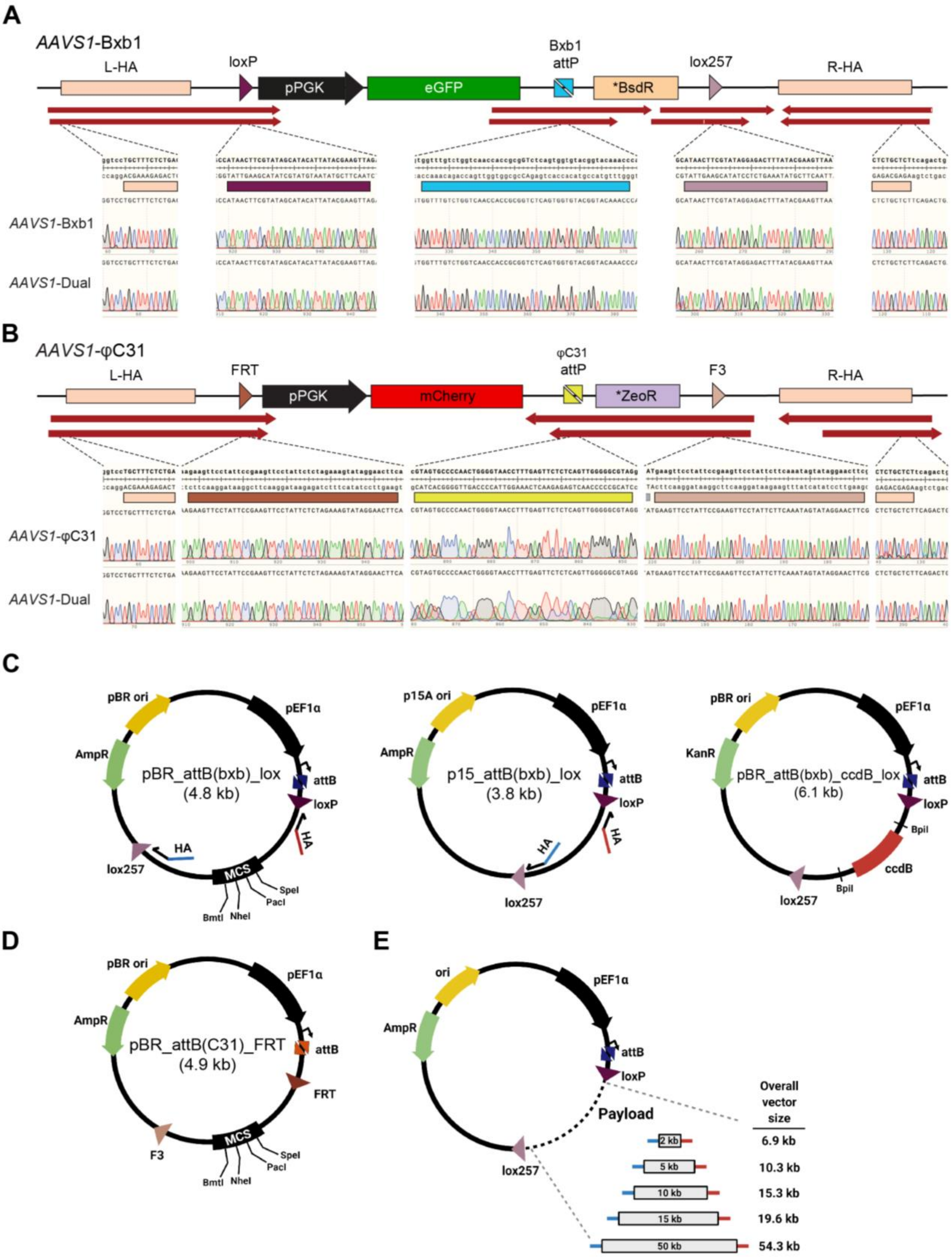
A
AAVS1-Bxb1
L-HA
loxP
pPGK
eGFP
Bxb1
attP
*BsdR
lox257
R-HA
AAVS1-Bxb1
AAVS1-Dual
B
AAVS1-φC31
L-HA
FRT
pPGK
mCherry
φC31
attP
*ZeoR
F3
R-HA
AAVS1-φC31
AAVS1-Dual
C
pBR ori
pEF1α
AmpR
pBR_attB(bxb)_lox
(4.8 kb)
attB
loxP
HA
lox257
MCS
SpeI
PacI
NheI
BmtI
p15A ori
p15_attB(bxb)_lox
(3.8 kb)
KanR
pBR_attB(bxb)_ccdB_lox
(6.1 kb)
BpiI
ccdB
D
pBR_attB(C31)_FRT
(4.9 kb)
FRT
F3
E
ori
Payload
Overall vector size
2 kb
5 kb
10 kb
15 kb
50 kb
6.9 kb
10.3 kb
15.3 kb
19.6 kb
54.3 kb

**Figure S1. Overview of the hiPSC *AAVS1*-acceptor lines and the donor vectors**

(A) Sanger sequencing confirming targeting of the Bxb1-LP to the *AAVS1* locus and the sequences of the *loxP, lox257* and *attP* (Bxb1) sites in the *AAVS1*-Bxb1 and *AAVS1*-Dual hiPSC lines. Red arrows indicate alignment of the sequencing chromatograms. L-HA, left homology arm; R-HA, right homology arm.

(B) Sanger sequencing confirming targeting of the φC31-LP to the *AAVS1* locus and the sequences of the *FRT*, *F3* and *attP* (φC31) sites in the *AAVS1*-φC31 and *AAVS1*-Dual hiPSC lines. Red arrows indicate alignment of the sequencing chromatograms. L-HA, left homology arm; R-HA, right homology arm.

(C) Schematics of the Bxb1 donor vectors used for inserting the DNA payloads and their respective sizes (kilobases, kb). The pBR_attB(bxb)_lox plasmid (*left*) was used for cloning payloads <20 kb, either by enzymatic digestion and ligation using restriction enzymes indicated within the multi-cloning site (MCS), or by recombineering. The p15_attB(bxb)_lox plasmid (*middle*) was used for cloning payloads between ~20-50 kb by recombineering. The pBR_attB(bxb)_ccdB_lox plasmid (*right*) was used for cloning payloads via modular assembly strategies involving digestion of the plasmid with the type IIS restriction enzyme, BpiI. Half arrows, recombineering primers used to amplify the cloning vector with homology arms (HA) to the DNA payload attached; pEF1a; human elongation factor 1 alpha promoter; ori, origin of replication; AmpR, b-lactamase; KanR, aminoglycoside phosphotransferase.

(D) Schematic of the φC31 donor vector (pBR_attB(C31)_FRT). pEF1a; human elongation factor 1 alpha promoter; ori, origin of replication; AmpR, b-lactamase.

(E) Overview of the Bxb1-donor vectors containing DNA payloads between ~2 – 50 kb, and the resulting size of the vector.

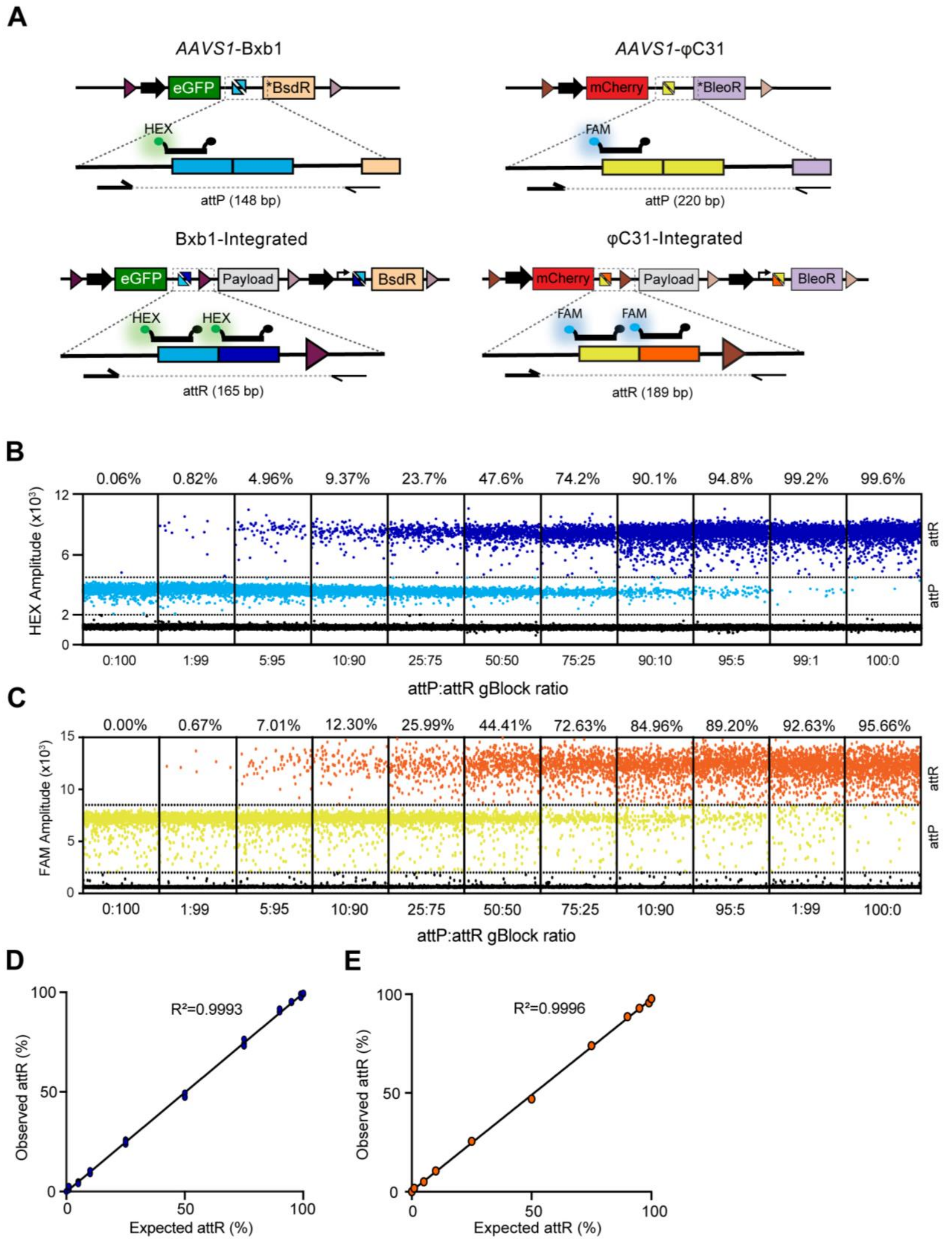


**Figure S2. Validation of *attP:attR* ddPCR assay for quantifying Bxb1- or φC31-mediated integration**

(A) Schematic of the DNA regions amplified from non-integrated (*attP*, *upper*) and integrated cells (*attR*, *lower*) in the ddPCR assay for either Bxb1 (*left*) or φC31 (*right*) mediated integration. A common forward primer (thick half arrow) and sequence-specific reverse primers (thin half arrows) were used to amplify the PCR products. Either one or both fluorescence probes (thick black bar) could anneal to the two resulting amplicons, leading to differences in signal intensity.

(B, C) Representative ddPCR dot plots of the observed frequencies at different ratios for two synthetic sequences matching the expected *attP* and *attR* amplicons for either Bxb1 (B) or φC31 (C). Sequences were spiked into genomic DNA. Dots represent droplets containing the indicated sequence, while percentages denote the calculated integration efficiency.

(D, E) Regression analysis for the observed versus expected frequency of the Bxb1 (D) and φC31 (E) *attR* amplicons; n=3 replicates.

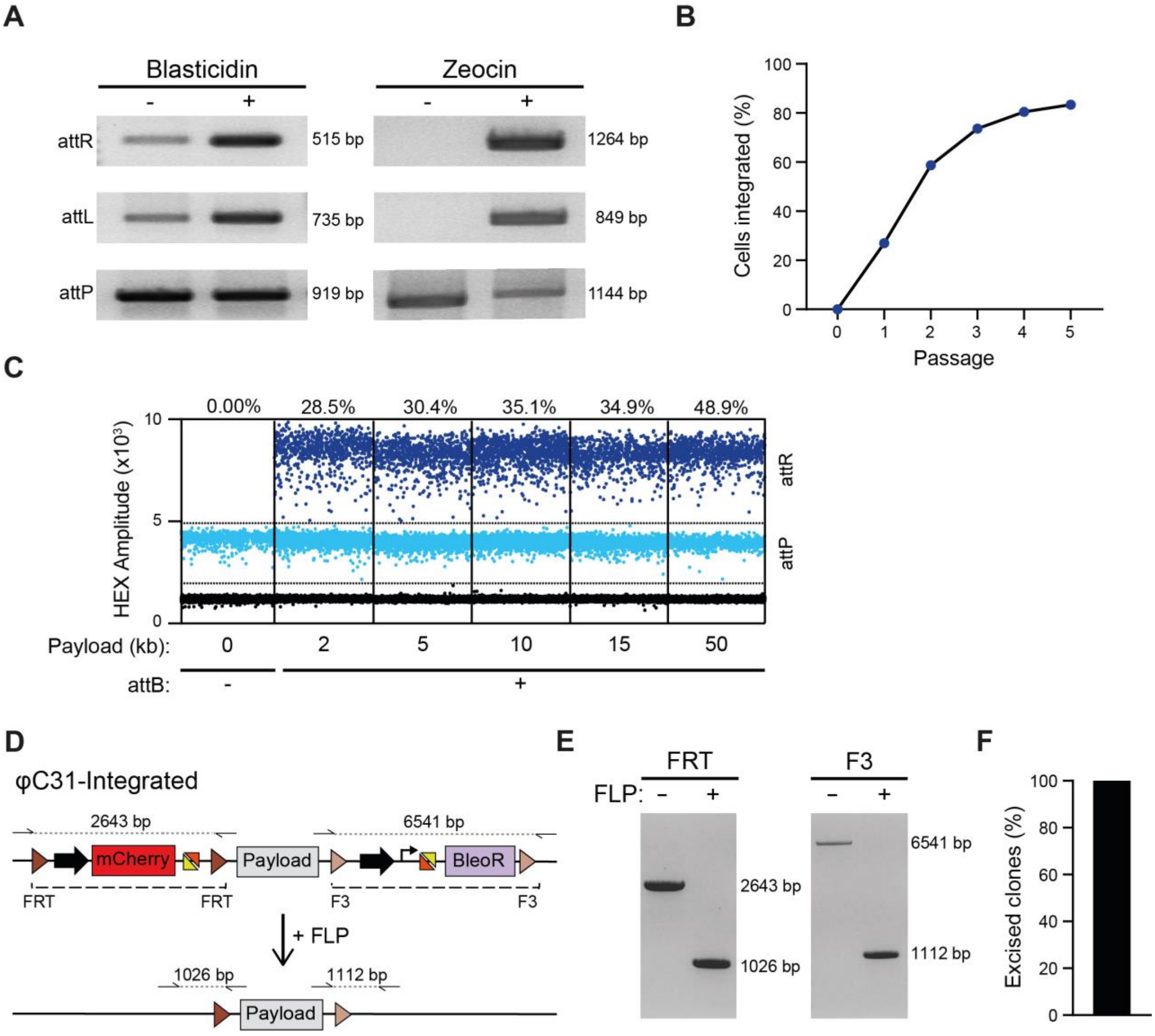


**Figure S3. Evaluating the efficiency of integration and excision using STRAIGHT-IN**

(A) PCR amplification of genomic DNA confirming integration of the Bxb1- and φC31-donor vectors into the *AAVS1*-Dual hiPSCs. The “-” and “+” symbols indicate before and after the corresponding antibiotic selection respectively.

(B) Percentage of cells with the Bxb1 donor vector integrated when blasticidin selection is maintained for 5 passages of the hiPSCs.

(C) Representative ddPCR dot plots of hiPSCs that have integrated donor constructs with or without the *attB* sequence (+ or -) and payloads ranging from 0 – 50 kb into the landing pad (LP) following one round of blasticidin selection. Dots represent droplets containing the indicated sequence (attR or attP), while percentages denote the calculated integration efficiency.

(D) Schematic of procedure for excising the positive selection cassettes and vector backbone following integration of the donor vector into the φC31-LP. Dashed lines indicate the sequences excised. Half arrows indicate primer binding sites with dotted lines representing the resulting PCR amplicons.

(E) PCR screening using primer pairs indicated in (D), confirming the reduction in amplicon length upon expression of FLP recombinase (+).

(F) Quantification of integrated hiPSCs that have excised the auxiliary sequences following FLP expression.

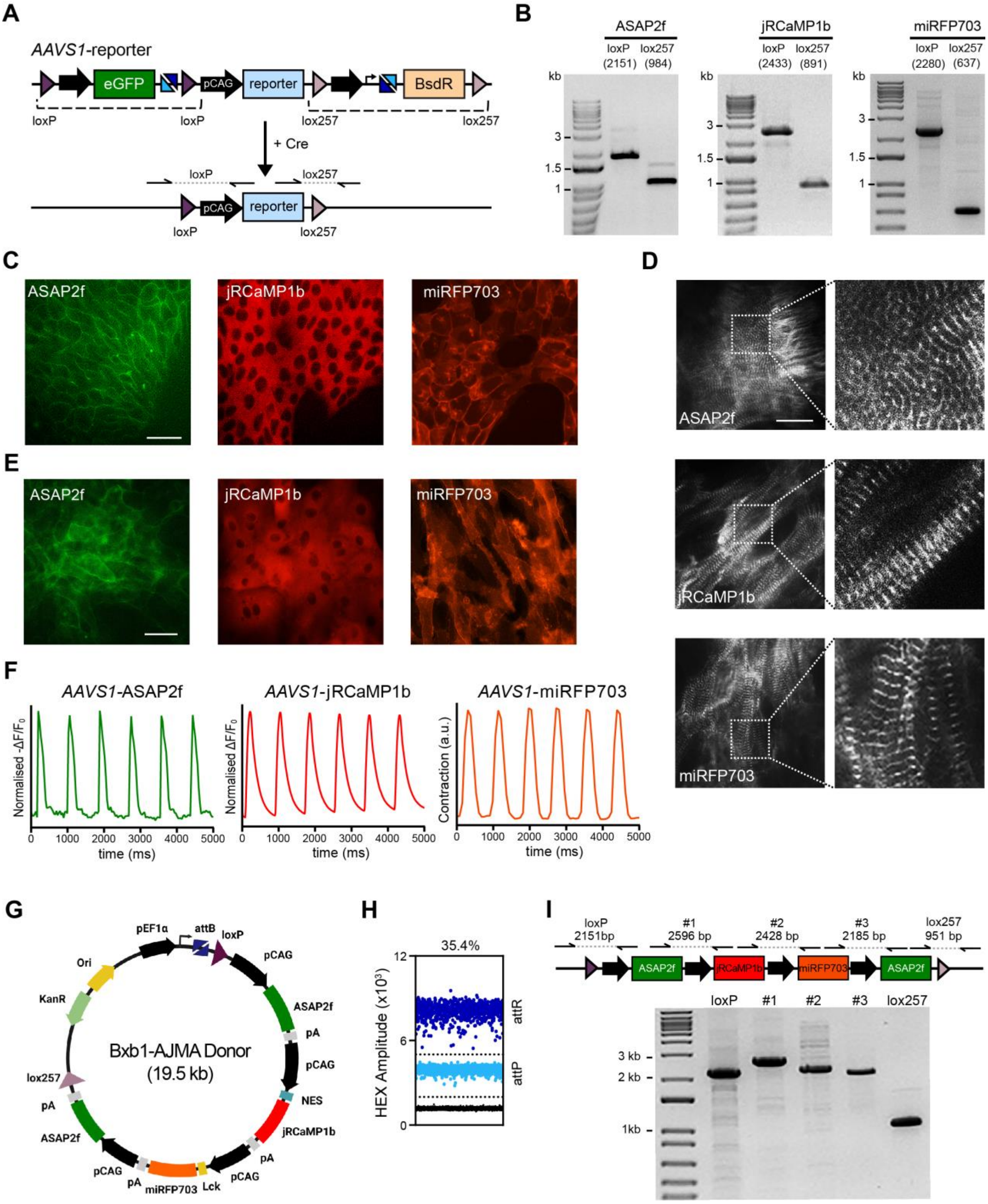


**Figure S4. Assessment of genetic reporters introduced by STRAIGHT-IN for evaluating APs, $Ca^{2+}$ transients and contractility kinetics in hiPSC-CMs**

(A) Schematic of the composition of the *AAVS1* locus following the integration of the fluorescent reporters (*ASAP2f, jRCaMP1b, miRFP703*). Filled black arrows, constitutive promoters; eGFP, enhanced green fluorescence protein; BsdR, blasticidin resistance gene; dashed lines, sequences excised; half arrows, primer binding sites; dotted lines, resulting PCR amplicons generated by screening across the remaining *loxP* and *lox257* sites.

(B) PCR screening across regions indicated in (A) confirming targeted integration of each of the fluorescent reporters into the *AAVS1*-Bxb1 hiPSC line. The base-pair size of the expected amplicons for each reporter is indicated in brackets. A DNA ladder was loaded in the first column of each gel, with the size of selected bands indicated. kb, kilobase.

(C) Fluorescence images from the *AAVS1*-reporter hiPSC lines indicating the cellular localisation of each of the integrated reporters. Scale bars, 50 µm.

(D) Immunofluorescence images of the cardiac sarcomeric protein α-actinin from each of the *AAVS1*-reporter hiPSC lines following differentiation to cardiomyocytes. Images on the right are magnifications of the regions within the dotted boxes. Scale bars, 25 µm.

(E) Fluorescence images of cardiomyocytes differentiated from the *AAVS1*-reporter hiPSC lines indicating the cellular localisation of each of the integrated reporters. Scale bars, 50 µm.

(F) Representative time plots of baseline-normalised fluorescence signals from the *AAVS1*-reporter hiPSC-CMs stimulated at 1.2 Hz. Changes in the fluorescence of *AAVS1*-ASAP2f (*left*) and *AAVS1*-jRCaMP1b (*middle*) hiPSC-CMs reflect the action potential and cytosolic $Ca^{2+}$ transients respectively, while the displacement of the fluorescence signal in *AAVS1*-miRFP703 hiPSC-CMs indicates contraction dynamics.

(G) Schematic of the donor vector for integrating two *ASAP2f* expression cassettes and single expression cassettes for *jRCaMP1b* and *miRFP703*. pEF1a; human elongation factor 1 alpha promoter; pCAG, CAG promoter; pA, polyadenylation signal; NES, nuclear export signal; Lck, Lck membrane targeting signal; ori, origin of replication; KanR, aminoglycoside phosphotransferase.

(H) ddPCR dot plot of *AAVS1*-Bxb1 hiPSCs transfected with the Bxb1-AJMA donor vector. Dots represent droplets containing the indicated sequence (attR or attP), while the percentage denotes the calculated integration efficiency.

(I) Schematic of the resulting *AAVS1*-AJMA locus (*top*). Half arrows, primer binding sites; dotted lines, resulting PCR amplicons with expected sizes indicated in brackets. PCR screening (*bottom*), using the primer pairs indicated, confirming targeted integration of the multi-reporter construct and no internal rearrangement of the transgenes. A DNA ladder was loaded in the first column of the gel, with the size of selected bands indicated. kb, kilobase.

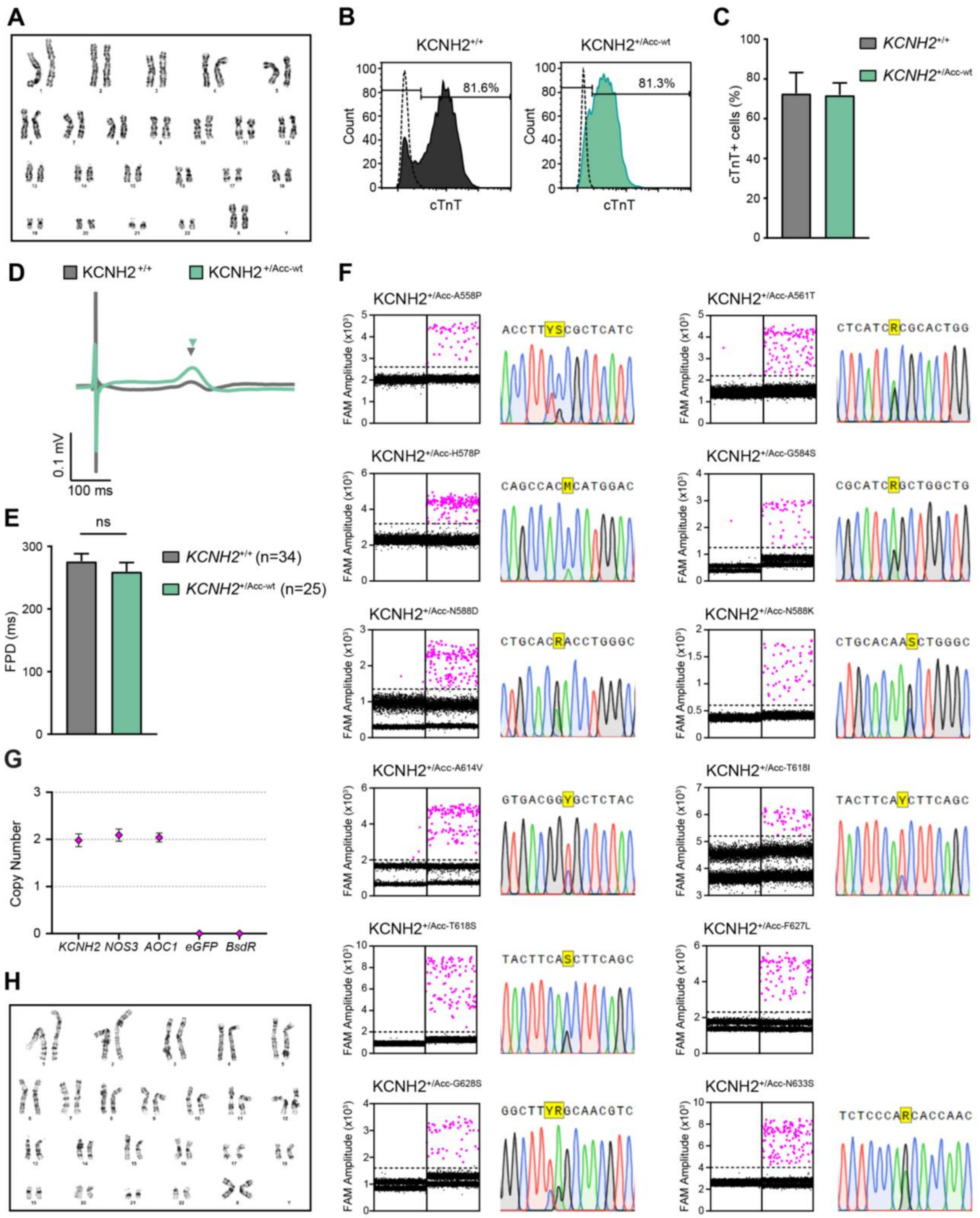


**Figure S5. Generating and characterising *KCNH2*-variant hiPSCs and hiPSC-CMs.**

(A) G-banding karyogram for a *KCNH2*$^{+/Acc\text{-}wt}$ hiPSC line generated by STRAIGHT-IN indicating a normal 46, XX karyotype.

(B) Representative histogram plots showing the percentage of cardiomyocytes (cTnT$^+$) as determined by flow cytometry following differentiation of either *KCNH2*$^{+/+}$ or *KCNH2*$^{+/Acc\text{-}wt}$ hiPSCs. Dotted lines represent a control cTnT$^-$ population.
(C) Averaged percentage of differentiated *KCNH2*$^{+/+}$ and *KCNH2*$^{+/Acc\text{-}wt}$ hiPSCs that stained positive for cTnT. n=3 differentiations and error bars represent ± SEM.
(D, E) Representative averaged field potential (FP) traces (D) and averaged FP duration (FPD) values (E) of *KCNH2*$^{+/+}$ and *KCNH2*$^{+/Acc\text{-}wt}$ hiPSC-CMs. Arrowheads indicate the repolarization peak for each trace. Values (n) indicate the number of recordings. Error bars represent ± SEM; ns, not significant ($p > 0.05$; unpaired t-test).
(F) Dot plots indicating the detection by ddPCR following Bxb1-mediated integration of the specified *KCNH2* variants (A558P; A561T; H578P; G584S; N588D; N588K; A614V; T618I; T618S; F627L; G628S; N633S) in a pool of transfected cells, with subsequent sequence analysis following Cre transfection and demultiplexing of the variants. Note, while the variant *KCNH2*$^{+/Acc\text{-}F627L}$ was integrated, clonal lines were not recovered hence why no chromatogram is available.
(G) ddPCR confirming that a *KCNH2*$^{+/Acc\text{-}A561T}$ hiPSC line generated by STRAIGHT-IN contained 2 copies of genomic genes *KCNH2*, *NOS3* and *AOC1*, and no copies of the Bxb1-LP cassette transgenes, *eGFP* and *BsdR.* Error bars represent Poisson 95% CI.
(H) G-banding karyogram for the *KCNH2*$^{+/Acc\text{-}A561T}$ hiPSC line indicating a normal 46, XX karyotype.

**Table S1. KCNH2 variants introduced in KCNH2$^{+/Acc}$ hiPSCs, annotated and with corresponding gBlock sequence**

| SNP | Nucleotide change | Protein change | Interpretation (ClinVar) | Associated Disease | gBlock | Comments |
|---|---|---|---|---|---|---|
| rs121912576 | c.1672G>C | A558P | pathogenic | Long QT syndrome | cttgccccccttgccccatcaacggaatgtgccccttccctgtcccccagctgatcgggctgctgaagactgcgcggctgctgcggctgg<br>tgcgcgtggcgcggaagctggatcgctactcagagtacggcgcggccgtgctgttcttgctcatgtgcaccttCCcgctcatcgcgcac<br>tggctagcctgcatctggtacgccatcggcaacatggagcagccacacatggactcacgcatcggctggctgcacaacctgggcgac<br>cagataggcaaaccctacaacagcagcggcctgggcggcccctccatcaaggacaagtatgtgacggcgctctacttcaccttcagc<br>agcctcaccagtgtgggcttcggcaacgtctctcccaacaccaactcagagaagatcttctccatctgcgtcatgctcattggctgtgag<br>tgtgcccaggggcgggcggcggggagagcccacggtggaggaacccaagttggaggaaactgaggctgctagccgggcca | silent mutation introduced into adjacent amino acid |
| rs199472921 | c.1681G>A | A561T | pathogenic | Long QT syndrome | cttgccccccttgccccatcaacggaatgtgccccttccctgtcccccagctgatcgggctgctgaagactgcgcggctgctgcggctgg<br>tgcgcgtggcgcggaagctggatcgctactcagagtacggcgcggccgtgctgttcttgctcatgtgcacctttgcgctcatcAcgcac<br>tggctagcctgcatctggtacgccatcggcaacatggagcagccacacatggactcacgcatcggctggctgcacaacctgggcgac<br>cagataggcaaaccctacaacagcagcggcctgggcggcccctccatcaaggacaagtatgtgacggcgctctacttcaccttcagc<br>agcctcaccagtgtgggcttcggcaacgtctctcccaacaccaactcagagaagatcttctccatctgcgtcatgctcattggctgtgag<br>tgtgcccaggggcgggcggcggggagagcccacggtggaggaacccaagttggaggaaactgaggctgctagccgggcca | |
| rs794728376 | c.1733 A>C | H578P | uncertain significance | - | cttgccccccttgccccatcaacggaatgtgccccttccctgtcccccagctgatcgggctgctgaagactgcgcggctgctgcggctgg<br>tgcgcgtggcgcggaagctggatcgctactcagagtacggcgcggccgtgctgttcttgctcatgtgcacctttgcgctcatcgcgcact<br>ggctagcctgcatctggtacgccatcggcaacatggagcagccacCcatggactcacgcatcggctggctgcacaacctgggcgacc<br>agataggcaaaccctacaacagcagcggcctgggcggcccctccatcaaggacaagtatgtgacggcgctctacttcaccttcagca<br>gcctcaccagtgtgggcttcggcaacgtctctcccaacaccaactcagagaagatcttctccatctgcgtcatgctcattggctgtgagt<br>gtgcccaggggcgggcggcggggagagcccacggtggaggaacccaagttggaggaaactgaggctgctagccgggcca | |
| rs199473428 | c.1750G>A | G584S | likely pathogenic; pathogenic | Long QT syndrome | cttgccccccttgccccatcaacggaatgtgccccttccctgtcccccagctgatcgggctgctgaagactgcgcggctgctgcggctgg<br>tgcgcgtggcgcggaagctggatcgctactcagagtacggcgcggccgtgctgttcttgctcatgtgcacctttgcgctcatcgcgcact<br>ggctagcctgcatctggtacgccatcggcaacatggagcagccacacatggactcacgcatcAgctggctgcacaacctgggcgacc<br>agataggcaaaccctacaacagcagcggcctgggcggcccctccatcaaggacaagtatgtgacggcgctctacttcaccttcagca<br>gcctcaccagtgtgggcttcggcaacgtctctcccaacaccaactcagagaagatcttctccatctgcgtcatgctcattggctgtgagt<br>gtgcccaggggcgggcggcggggagagcccacggtggaggaacccaagttggaggaaactgaggctgctagccgggcca | |
| rs199473431 | c.1762A>G | N588D | not provided | Long QT syndrome | cttgccccccttgccccatcaacggaatgtgccccttccctgtcccccagctgatcgggctgctgaagactgcgcggctgctgcggctgg<br>tgcgcgtggcgcggaagctggatcgctactcagagtacggcgcggccgtgctgttcttgctcatgtgcacctttgcgctcatcgcgcact<br>ggctagcctgcatctggtacgccatcggcaacatggagcagccacacatggactcacgcatcggctggctgcacGacctgggcgacc<br>agataggcaaaccctacaacagcagcggcctgggcggcccctccatcaaggacaagtatgtgacggcgctctacttcaccttcagca<br>gcctcaccagtgtgggcttcggcaacgtctctcccaacaccaactcagagaagatcttctccatctgcgtcatgctcattggctgtgagt<br>gtgcccaggggcgggcggcggggagagcccacggtggaggaacccaagttggaggaaactgaggctgctagccgggcca | |
| rs104894021 | c.1764C>G | N588K | pathogenic | Short QT syndrome | cttgccccccttgccccatcaacggaatgtgccccttccctgtcccccagctgatcgggctgctgaagactgcgcggctgctgcggctgg<br>tgcgcgtggcgcggaagctggatcgctactcagagtacggcgcggccgtgctgttcttgctcatgtgcacctttgcgctcatcgcgcact<br>ggctagcctgcatctggtacgccatcggcaacatggagcagccacacatggactcacgcatcggctggctgcacaaGctgggcgacc<br>agataggcaaaccctacaacagcagcggcctgggcggcccctccatcaaggacaagtatgtgacggcgctctacttcaccttcagca<br>gcctcaccagtgtgggcttcggcaacgtctctcccaacaccaactcagagaagatcttctccatctgcgtcatgctcattggctgtgagt<br>gtgcccaggggcgggcggcggggagagcccacggtggaggaacccaagttggaggaaactgaggctgctagccgggcca | |

**Table S1** *(continued)*

| SNP | Nucleotide change | Protein change | Interpretation (ClinVar) | Associated Disease | gBlock | Comments |
|---|---|---|---|---|---|---|
| rs199472944 | c.1841C>T | A614V | pathogenic | Long QT syndrome | cttgccccccttgccccatcaacggaatgtgccccttccctgtcccccagctgatcgggctgctgaagactgcgcggctgctgcggctgg<br>tgcgcgtggcgcggaagctggatcgctactcagagtacggcgcggccgtgctgttcttgctcatgtgcacctttgcgctcatcgcgcact<br>ggctagcctgcatctggtacgccatcggcaacatggagcagccacacatggactcacgcatcggctggctgcacaacctgggcgacc<br>agataggcaaaccctacaacagcagcggcctgggcggcccctccatcaaggacaagtatgtgacggTgctctacttcaccttcagca<br>gcctcaccagtgtgggcttcggcaacgtctctcccaacaccaactcagagaagatcttctccatctgcgtcatgctcattggctgtgagt<br>gtgcccaggggcgggcggcggggagagcccacggtggaggaacccaagttggaggaaactgaggctgctagccgggcca | |
| rs199472947 | c.1853C>T | T618I | not provided | Short QT syndrome | cttgccccccttgccccatcaacggaatgtgccccttccctgtcccccagctgatcgggctgctgaagactgcgcggctgctgcggctgg<br>tgcgcgtggcgcggaagctggatcgctactcagagtacggcgcggccgtgctgttcttgctcatgtgcacctttgcgctcatcgcgcact<br>ggctagcctgcatctggtacgccatcggcaacatggagcagccacacatggactcacgcatcggctggctgcacaacctgggcgacc<br>agataggcaaaccctacaacagcagcggcctgggcggcccctccatcaaggacaagtatgtgacggcgctctacttcaTcttcagca<br>gcctcaccagtgtgggcttcggcaacgtctctcccaacaccaactcagagaagatcttctccatctgcgtcatgctcattggctgtgagt<br>gtgcccaggggcgggcggcggggagagcccacggtggaggaacccaagttggaggaaactgaggctgctagccgggcca | |
| rs199472947 | c.1853C>G | T618S | not provided | Long QT syndrome | cttgccccccttgccccatcaacggaatgtgccccttccctgtcccccagctgatcgggctgctgaagactgcgcggctgctgcggctgg<br>tgcgcgtggcgcggaagctggatcgctactcagagtacggcgcggccgtgctgttcttgctcatgtgcacctttgcgctcatcgcgcact<br>ggctagcctgcatctggtacgccatcggcaacatggagcagccacacatggactcacgcatcggctggctgcacaacctgggcgacc<br>agataggcaaaccctacaacagcagcggcctgggcggcccctccatcaaggacaagtatgtgacggcgctctacttcaGcttcagca<br>gcctcaccagtgtgggcttcggcaacgtctctcccaacaccaactcagagaagatcttctccatctgcgtcatgctcattggctgtgagt<br>gtgcccaggggcgggcggcggggagagcccacggtggaggaacccaagttggaggaaactgaggctgctagccgggcca | |
| rs199473039 | c.1881C>A/G | F627L | pathogenic | Long QT syndrome | cttgccccccttgccccatcaacggaatgtgccccttccctgtcccccagctgatcgggctgctgaagactgcgcggctgctgcggctgg<br>tgcgcgtggcgcggaagctggatcgctactcagagtacggcgcggccgtgctgttcttgctcatgtgcacctttgcgctcatcgcgcact<br>ggctagcctgcatctggtacgccatcggcaacatggagcagccacacatggactcacgcatcggctggctgcacaacctgggcgacc<br>agataggcaaaccctacaacagcagcggcctgggcggcccctccatcaaggacaagtatgtgacggcgctctacttcaccttcagca<br>gcctcaccagtgtgggcttGggcaacgtctctcccaacaccaactcagagaagatcttctccatctgcgtcatgctcattggctgtgag<br>tgtgcccaggggcgggcggcggggagagcccacggtggaggaacccaagttggaggaaactgaggctgctagccgggcca | |
| rs121912507 | c.1882G>A | G628S | pathogenic | Long QT syndrome | cttgccccccttgccccatcaacggaatgtgccccttccctgtcccccagctgatcgggctgctgaagactgcgcggctgctgcggctgg<br>tgcgcgtggcgcggaagctggatcgctactcagagtacggcgcggccgtgctgttcttgctcatgtgcacctttgcgctcatcgcgcact<br>ggctagcctgcatctggtacgccatcggcaacatggagcagccacacatggactcacgcatcggctggctgcacaacctgggcgacc<br>agataggcaaaccctacaacagcagcggcctgggcggcccctccatcaaggacaagtatgtgacggcgctctacttcaccttcagca<br>gcctcaccagtgtgggcttTAgcaacgtctctcccaacaccaactcagagaagatcttctccatctgcgtcatgctcattggctgtgagt<br>gtgcccaggggcgggcggcggggagagcccacggtggaggaacccaagttggaggaaactgaggctgctagccgggcca | silent mutation introduced into adjacent amino acid |
| rs199472961 | c.1898A>G | N633S | pathogenic | Long QT syndrome | cttgccccccttgccccatcaacggaatgtgccccttccctgtcccccagctgatcgggctgctgaagactgcgcggctgctgcggctgg<br>tgcgcgtggcgcggaagctggatcgctactcagagtacggcgcggccgtgctgttcttgctcatgtgcacctttgcgctcatcgcgcact<br>ggctagcctgcatctggtacgccatcggcaacatggagcagccacacatggactcacgcatcggctggctgcacaacctgggcgacc<br>agataggcaaaccctacaacagcagcggcctgggcggcccctccatcaaggacaagtatgtgacggcgctctacttcaccttcagca<br>gcctcaccagtgtgggcttcggcaacgtctctcccaGcaccaactcagagaagatcttctccatctgcgtcatgctcattggctgtgagt<br>gtgcccaggggcgggcggcggggagagcccacggtggaggaacccaagttggaggaaactgaggctgctagccgggcca | |

*Introduced mutations are indicated in capital letters.*
